# Inferring wildlife poaching in Southeast Asia with multispecies dynamic occupancy models

**DOI:** 10.1101/520114

**Authors:** Lucile Marescot, Arnaud Lyet, Rohit Singh, Neil Carter, Olivier Gimenez

## Abstract

Determining the ‘space race’ between sympatric species is crucial to understand the effects of interspecific interactions on the extinction risk of species threatened by poachers, predators, pathogens, and invasive competitors. Dynamic two-species occupancy models provide a flexible framework to decompose complex species interaction patterns while accounting for imperfect detection. In particular, these models can describe poachers-wildlife interactions by considering the occupancy, the extinction and colonisation probabilities of wildlife conditional on the presence or absence of poachers and vice versa. We apply our model to a case study on wildlife poaching in the Eastern plains of Cambodia. We used co-occurrence data extracted from the database of the SMART partnership to study the distribution dynamics between poachers and six ungulate species regarded as main prey of tigers. We used 4 years of survey data reporting the locations of ranger patrols on the detection of snares with visual detections or presence signs of the ungulates. Our results showed that a substantial proportion of the sites occupied by ungulate species went extinct over the years of the study while the proportion of sites colonised by poachers increased. We also showed, for the first time, that spatio-temporal heterogeneity in the patrolling effort explains a great deal of the variation in the detection of poachers and ungulates. Our approach provides practitioners with a flexible and robust tool to assess conservation status of species and extinction risk of wildlife populations. It can assist managers in better evaluating, learning and adapting the patrolling strategies of rangers.

## Introduction

Illegal hunting (hereafter poaching) is a major threat to wildlife species around the world (Gray et al. 2018; Ripple et al. 2016). Poaching of endangered wildlife, in particular, can push those species to the brink of extinction (Gross 2018). Moreover, poaching of certain taxa, like ungulates, can deplete prey resources of large carnivore species. Prey depletion is considered a driving force of global declines in carnivore numbers (Wolf & Ripple 2016). Thus, quantifying poaching pressure in both space and time is an urgent conservation need. Doing so would help causally link poaching behaviour to wildlife population dynamics and better inform intervention efforts. Yet, assessing the interrelationships between poachers and wildlife is notoriously difficult for various reasons, including: 1) inadequate information on the mediating influence of ranger patrols on poacher-wildlife interactions; 2) lack of empirical data on poachers, wildlife, and patrols at sufficient spatial and temporal resolutions; 3) imperfect detection of poachers and wildlife; and 4) the absence of statistical models to partition the influence of poachers, patrols, and wildlife on each other in space and time. To help overcome these difficulties, we build upon a rapidly advancing class of occupancy models to study spatiotemporal interactions between poachers and threatened wildlife species in Cambodia. This study is part of a larger effort to use ecological theory and models to inform wildlife restoration efforts in this region, including the reintroduction of the globally endangered tiger *(Panthera tigris)*.

Poachers and wildlife interactions can be compared to a predator-prey system in which interactions fluctuate over time in a “landscape of fear” (Laundre et al. 2010). Predator-prey interactions often represent a “space race”, where prey try to minimize and predators maximize spatial overlap (Sih 2005). Most empirical studies focusing on the “space race” for predator-prey interactions generally use telemetry or GPS tracking devices, assuming perfect detection probability of both predators and prey (Cusack et al. 2019). Tracking poachers with GPS devices is infeasible, so rangers often use indirect methods based on observations of snares, hunting camps, and social surveys to detect their presence (Milner-Gulland & Leader-Williams 1992). However, these observations suffer from imperfect detection of poachers and could therefore generate inaccurate estimates of poaching pressure or changes in wildlife populations, as well as lead to ineffective conservation policies.

Over the last 15 years, the occupancy modelling framework has been proposed and extended to include multiple interacting species to address the issues of imperfect species detection and dynamic changes in species distributions (MacKenzie et al. 2006, MacKenzie et al. 2018). As such, this framework is well suited to study spatiotemporal interactions between wildlife and poachers. Using repeated surveys of detections and non-detections of poaching-related threats, such as snares, and of wildlife collected at several spatial units on repeated occasions over several years, one can estimate the temporal variation in the detection and occupancy probabilities of poachers in a given site conditional on the occurrence of wildlife with which they interact, thereby simultaneously assessing the dynamics of interactions (Fidino et al. 2018). However, few studies account for the imperfect detection in the occupancy of poachers (Critchlow et al. 2015, Moore et al. 2018). One recent study assessed levels of illegal wildlife killing in a national park in Rwanda by fitting an occupancy model on detection/non-detection of poaching signs directly (Moore et al. 2018). This study did not account explicitly for interactions between poachers and wildlife. To our knowledge, no study has quantified how poachers and wildlife affect the space use of each other dynamically over time, while accounting for imperfect detection of both wildlife and poachers.

Here, for the first time, we utilized a two-species, multi-season occupancy model to assess the spatial dynamics of local interactions between poachers and common prey species of the tiger. According to conservationists, the tiger is functionally extinct in Cambodia since 2016, because of direct poaching and poaching of its main prey (CA|TS manual 2018). Our study provides a quantification of the impact of poaching on the occupancy of main tiger prey during the 4 years preceding tigers’ extinction from the region. Our study benefits from detection/non-detection data that were collected from January 2013 to December 2016 and georeferenced by rangers looking for poacher snares and signs of wildlife (e.g., hair, scat, footprint) during their patrols. As no records of tigers were found during those years, we only consider occurrences of six species regarded as the main tiger prey: the banteng, the gaur *(Bos gaunis)*, the sambar deer *(Rusa unicolor)*, the eld’s deer *(Rucervus eldii)* and two least-concern species, the wild boar *(Sus scrofa)* and the barking deer, also called red muntjac *(Muntiacus muntjak)* (Gray and Phan 2011).

Our model allows us to 1) investigate effects of spatial and temporal patterns in patrolling effort on detection and occupancy of poachers and wildlife; 2) estimate site-specific extinction/colonisation probabilities of wildlife conditional on the presence of poachers; and 3) estimate the extinction/colonisation probability of poachers conditional on the presence of wildlife. Thus, we have developed the two following hypotheses. First, poachers drive the dynamic space race despite rangers’ patrolling efforts (the “poachers winning” hypothesis). We therefore predict an increase in poachers’ occupancy and a decrease in wildlife occupancy. Alternatively, wildlife wins the space race despite the impact of poaching. In the “wildlife winning” hypothesis, we predict an increase or a stabilization of wildlife occupancy and a decrease in poachers’ occupancy as a function of patrolling effort (other hypotheses are described in the methods and appendix A). We need a better understanding of how poachers affect wildlife at fine spatial scales (e.g., within protected area borders). Our study helps do this by leveraging ecological theory and rapidly advancing statistical models. It also helps bridge the gap between science-based wildlife management and law enforcement to better inform ranger patrol strategies (Hofer et al. 2000, Hillborn et al. 2006).

## Material and methods

### Study area

We conducted our study in two protected areas within the eastern Plains Landscape of Cambodia: the Phnom Prich Wildlife Sanctuary (PPWS), covering more than 2,000 km^2^, and the Serepok Wildlife Sancturary (SWS), covering over 3,700 km of the Lower Mekong Dry Forest Eco-region. Both sites are among the 200 most important eco-regions of the world due to the high level of biodiversity facing myriad threats (Gray and Phan 2011).

Combined, the sanctuaries had total 11 patrol stations (5 in PPWS & 6 in SWS) from which rangers departed by motorbike (95% of the data), boats or vehicle, in order to inspect for illegal activities and monitor wildlife. Patrolling for illegal activity mainly consisted of locating, removing snares, intercepting poacher’s snares, and removing them. It also aimed at systematically recording any wildlife sign they may find during their patrolling activities.

### Data collection

The data we used for the analysis were obtained from the Spatial Monitoring And Reporting Tool (SMART), a tool that has been developed by the consortium of conservation organizations to measure, evaluate and improve the effectiveness of wildlife law enforcement patrols and site-based conservation activities (smartconservationtoools.org). SMART is also a key tool for the recovery of the populations of tiger and other keystone species (CA|TS manual 2018).

We extracted all the dates and geolocations of information collected by rangers from January 2013 to December 2016. We gathered 1647 GPS waypoints reported by rangers associated with 11 different patrolling stations. During each patrolling session, rangers recorded several observations on specific landscape features (e.g., roads and rivers), illegal activity (e.g., snaring, logging, fishing) and wildlife (direct sightings or signs of animal presence) or when no activity was detected.

For the wildlife data in the model, we used observations records and presence signs of the six tiger prey species mentioned above. We assumed rangers identified the species correctly during their field survey and if a misidentification occurred, it would be among the six ungulate species, which we pooled into one guild. We removed from the data the observations of carcasses that can be displaced by humans or other animals over long distance and therefore do not have reliable location. We used snares as relevant indicators of the presence of poachers and of their interactions with wildlife, as they represented a non-targeted catching technique for small and large wildlife species. Rangers collected the date and the GPS location of snares during their anti-poaching patrol route. Hereafter, we used the term “site occupied by poachers” to refer to the sites that have snares.

We plotted all the GPS data points collected in the two study areas, and used 10 x 10 km cells to partition the study areas, resulting in 42 sites for PPWS and 56 sites for SWS (Figure 1). We chose this cell size to represent sample sites in order to achieve a compromise between ecological and monitoring assumptions. We made sure the site was large enough to meet the closure assumption upon which occupancy models rely on (see section below), meaning that wildlife would not move out of the site between sampling occasions. On the other hand, we ensured the scale of the cells was sufficiently fine to ensure that rangers patrolled a fair proportion of each cell at least once during a primary occasion (Appendix B). Finally, we combined information on poaching-related threats and wildlife in a table composed of 81 rows representing the sites prospected by rangers and 48 columns representing the monthly occasions during the four years of the study period. We removed 17 sites never patrolled by rangers (Figure 1); seven cells were from the PPWS grid and ten from the SPS one.

**Figure 1:**
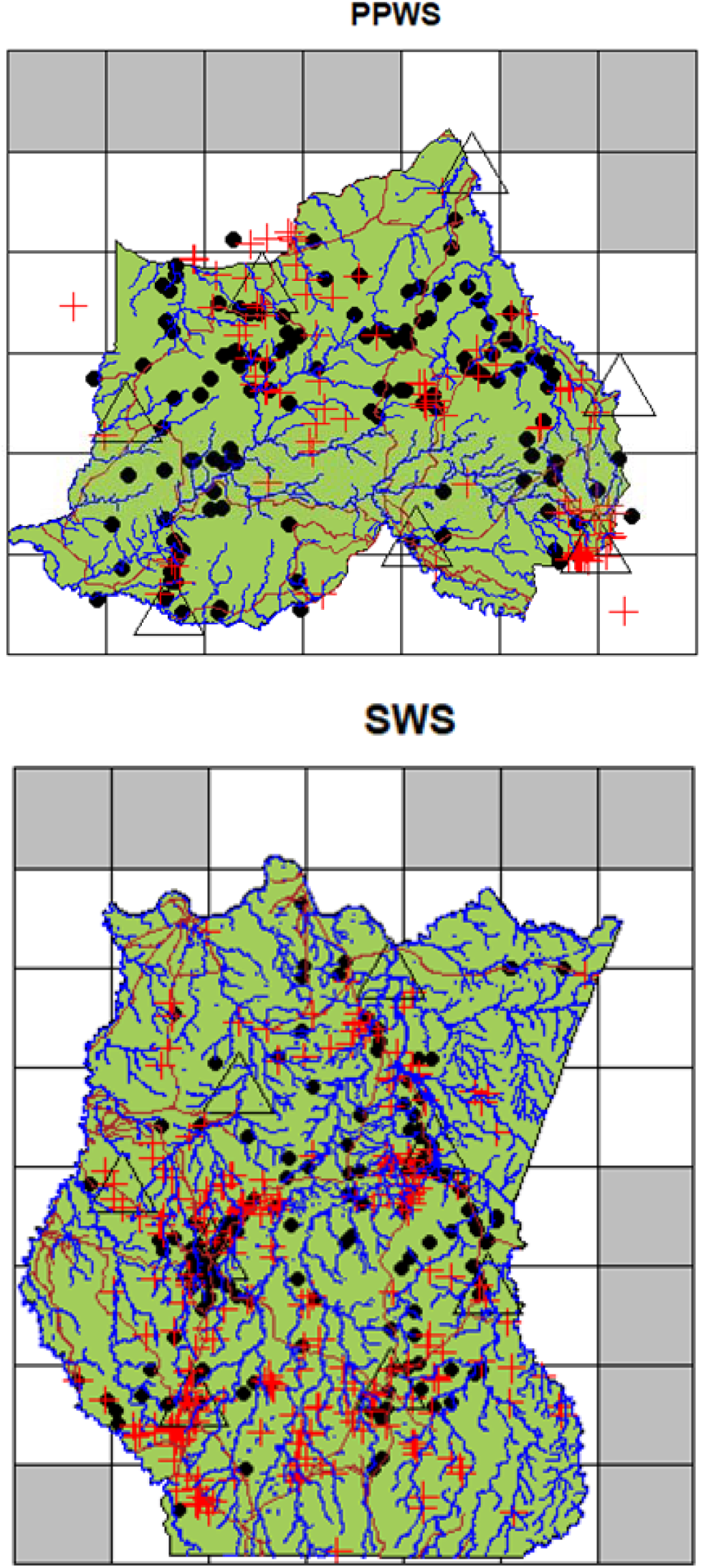
Map of the study areas in Cambodia, the Phnom Prich Wildlife Sanctuary (PPWS) and Serepok Wildlife Sanctuary (SWS) within the green polygons. Sampling sites were defined as 10×10 km cells. We show the occurrence patterns of poachers (red crosses) and wildlife species (black dots). Brown lines represent the main patrolling path network (roads), blue lines describe the river streams, black triangles represent the ranger stations as well as temporary ranger’s campsites. Note that only 81 cells were used for the analysis as we removed 17 sites that were never sampled (cells shaded in grey).

### Hidden Markov formulation of multi-season two-species occupancy model

#### Definition of states and observations

We developed a multi-season two-species occupancy model (MacKenzie et al. 2004) in a hidden Markov modelling (HMM) framework (Gimenez et al. 2014, Fiske et al. 2014). We defined the ecological process as a latent variable indicating at each sampling occasion whether a site was: “unoccupied by poachers and wildlife” (U), “occupied by poachers only” (OP), “occupied by wildlife only” (OW), and “occupied by wildlife and poachers” (WP). Occupancy models rely on the main assumption known as “population closure”, whereby sites are supposed to remain in the same occupancy states and only detection can vary between repeated surveys called secondary occasions. Sites can then change from one state to another between successive primary occasions through extinction and colonisation processes. The observation process was characterized by a known variable showing time series of detection/non-detection of species in each site (MacKenzie et al. 2006).

We adopted the Richmond-Waddle (RW) parameterization for occupancy and detection because it is more appropriate to describe poacher-wildlife interactions and identify the effects of covariates described below on species occupancy (Waddle et al. 2010, Richmond et al. 2010). This parameterization states that the initial occurrence of one species depends on the occurrence of the common species but the reciprocate regarding the common species is not true. We used the MacKenzie et al. (2006) parameterization for the transition process from one state to another, a convenient approach to model reciprocal responses between species following their interactions, meaning that extinction and colonisation probabilities of a species are conditional on the presence or absence of the other. This parameterization is important to predict between poachers and wildlife, which of the two, wins the ‘space race’ explained by underlying ecological mechanisms. It also considered the probability that a species replaced another in a given site from one primary occasion to another (Fidino et al. 2018).

#### Occupancy

To build the model, we first defined whether a site was initially occupied or not, and in which state, which was conveniently captured by the initial state probabilities of HMM (Gimenez et al. 2014). We proceed in two steps for clarity. The first step represents whether a site is occupied or not by wildlife regardless of the presence or absence of poachers. The following vector shows the probability of being in an unoccupied site U or a site occupied by W:

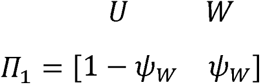

with *ψ_P/W_* the occupancy probability by wildlife and its complementary the probability of a site being unoccupied. The second matrix represents whether poachers, depending on the presence or absence of wildlife, occupy a site:

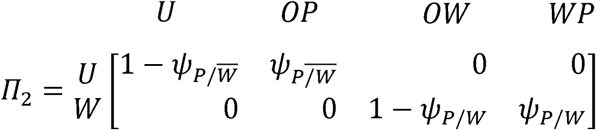

with *ψ_P/W_* the occupancy probability by poachers, conditional on the presence of wildlife and the complementary 1 − *ψ_P/W_* that a site is not occupied by poachers, conditional on the presence of wildlife. Parameter 1 − *ψ_P/W̄_* is the occupancy probability by poachers, conditional on the absence of wildlife and the complementary 1 − *ψ_P/W̄_* is the probability that poachers do not occupy a site conditional on the absence of animals.

#### Transitions

Conditional on the initial occupancy state, we describe how the state at a site changes over time assuming a Markovian process. We define the transition matrix *T* for a given site between the states U, OP, OW, WP from primary occasion *t* to the next primary occasion *t* + 1 as follows: 

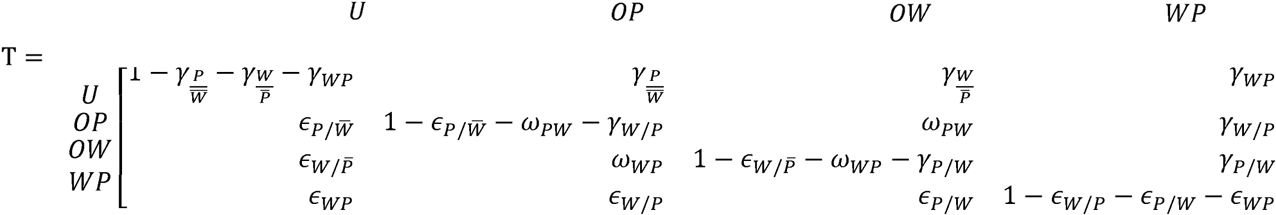

Each element of the matrix *T* is the transition probability from one of the occupancy states during a given year to another or the same occupancy state in the next year. The matrix describes four main processes: colonisation, extinction, replacement, and fidelity (MacKenzie 2018) that are detailed in Table 1 in appendix 1.

**Table 1:**
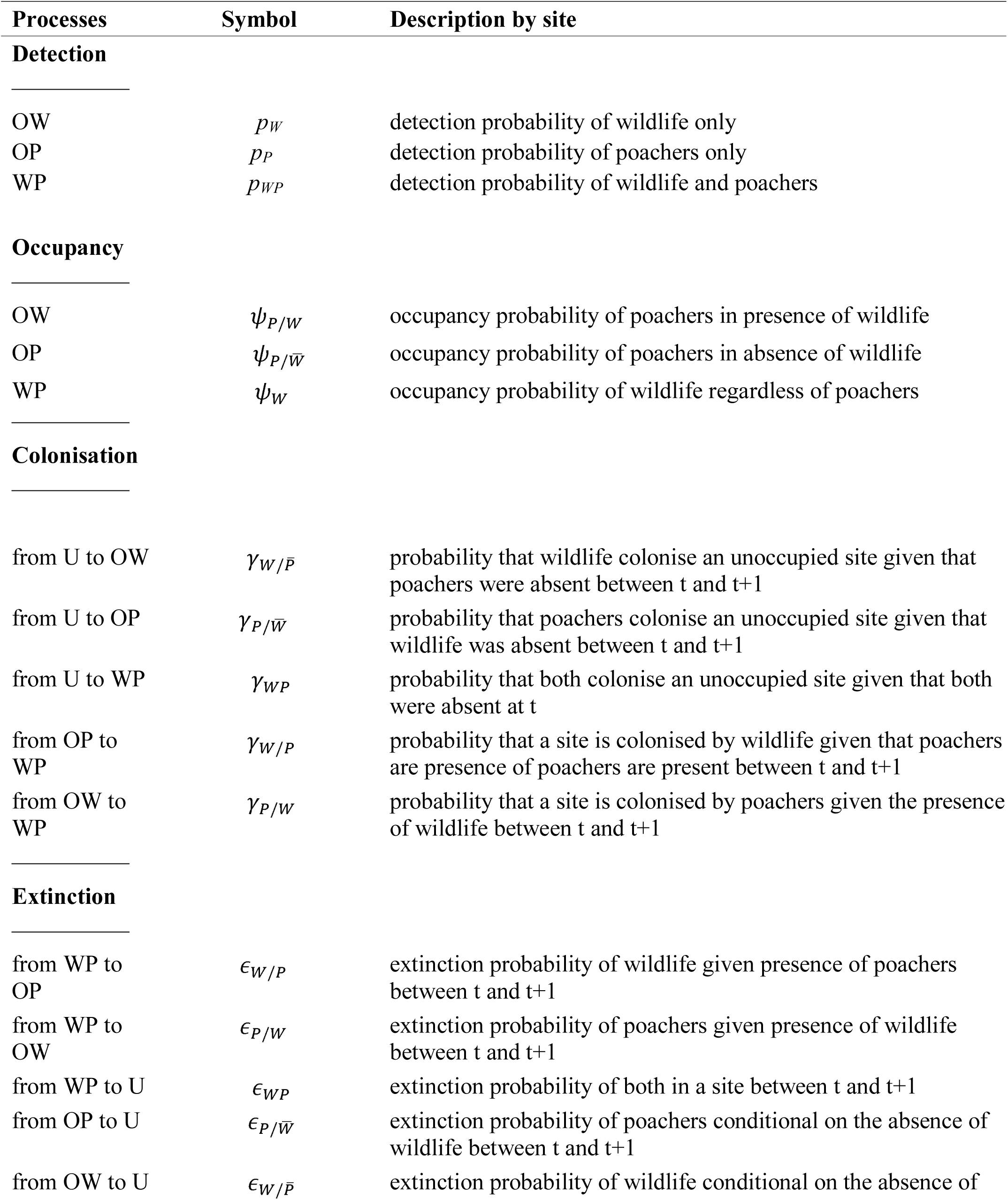
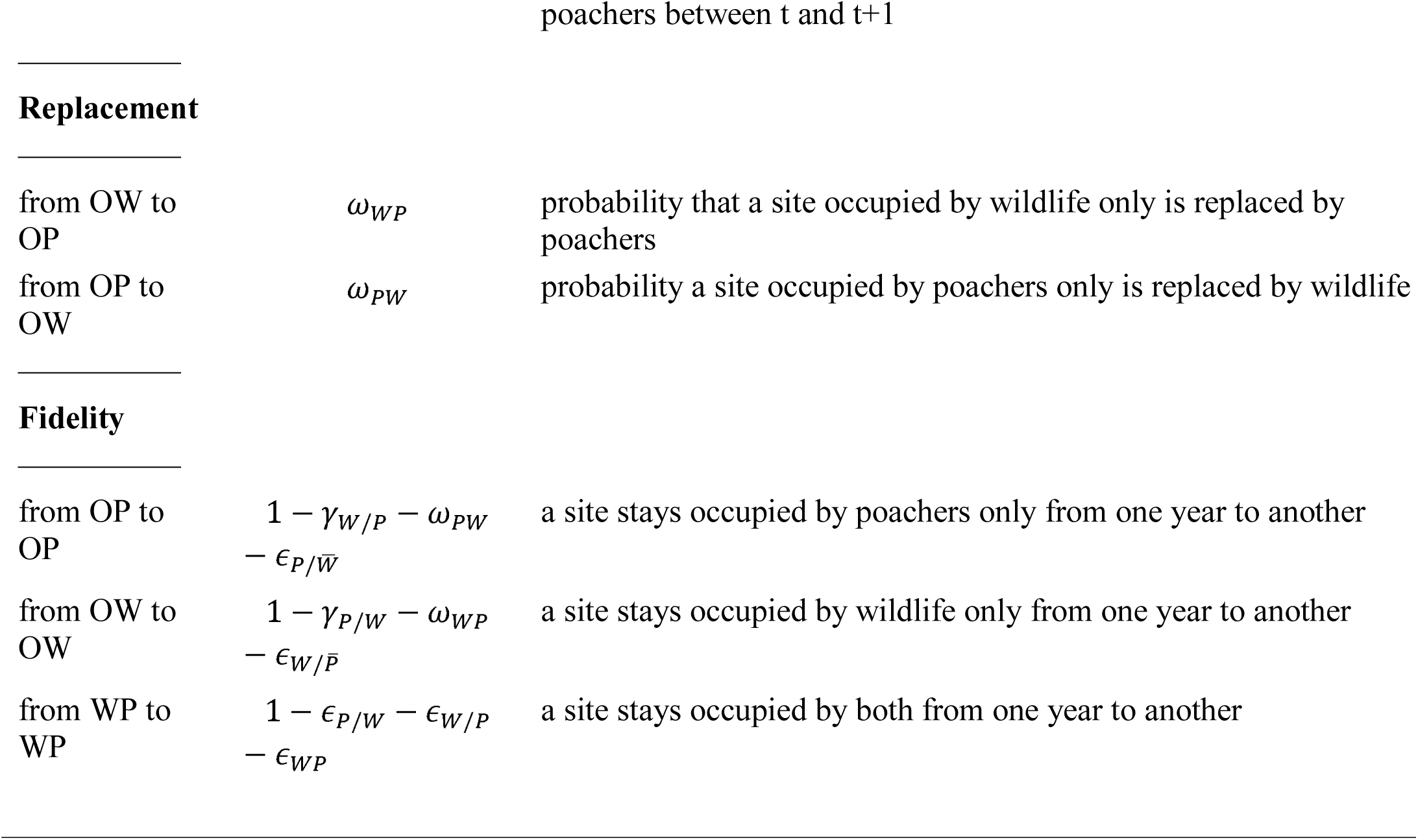
Description of the parameters used in the multi-season two-species occupancy model of dynamic interactions between poachers and wildlife.

These transitions occur at the beginning of each year (January). Within each primary occasion, the transition matrix *T* is simply a diagonal matrix of 1’s because the status of a site does not change from one secondary occasion to the other according to the closure assumption. Here, we defined the secondary occasions on a monthly-based time interval (from February to December).

#### Observations

The last step of the HMM consists of linking the observation process to the partially observed latent states (U, OW, OP, WP) describing the dynamic of the species occupancy. Four events represent the observation process: nothing was detected (ND), only wildlife was detected (WD), only poachers were detected (PD), or both were detected (WPD). For the sake of simplicity, we described detection probabilities of a species (poachers or wildlife) not conditionally on the presence/absence of the other (Fidino et al. 2018). We therefore defined the observation matrix as: 

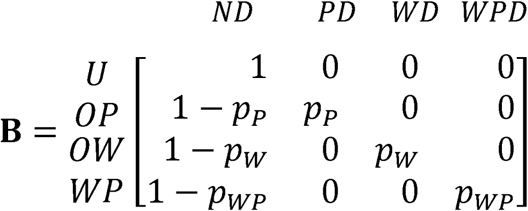

with *p_P_* the probability of detecting poachers in a site given that only poaching-related threats occur, *p_W_*the probability of detecting wildlife in a site where only wildlife occurs and *p_WP_* the probability of detecting poachers and wildlife conditional on the presence of both.

## Analyses

### Covariates

We accounted for the patrolling effort defined as the number of surveys carried out on each site during each sampling occasion and standardized to a scale from 0 when a site was not patrolled on a given occasion, to 1 where the most GPS waypoints were recorded on a sampling occasion (Appendix B). We also considered the nearest distance to a ranger station as another site-specific covariate reflecting spatial variation in the patrolling effort. We defined the shortest distance of each site to a road to account for effects of road access on the monitoring effort of rangers but also on the distributions of wildlife and poachers. Finally, we assessed the stream length crossing in each cell as another potential determinant of species detection and occupancy (Plumptre et al. 2010).

### Model selection and parameter estimation

We performed model selection on the poacher-wildlife occupancy and tested different hypotheses to determine the biotic (species interactions) and abiotic drivers (effects of roads, rivers, patrolling station, patrolling effort) of species detection, occupancy and transition probabilities. This was done in three steps.

First, we considered detection probability to differ between states of site occupation and varied with environmental covariates (see appendix A for model formulation). We used the best model structure on detection, and started the model selection procedure with a consensus model assuming a two-way interaction between poachers and wildlife, where occupancy and transition parameters of a species (wildlife or poachers) depend on whether the other was present. Then, we tested other ecological hypotheses, specific to poachers’ responses to wildlife and rangers and other environmental covariates. We looked at whether poachers’ initial occupancy was independent on the presence of wildlife but conditional of patrolling effort, or was independent on the presence of rangers but conditional to the presence of wildlife or independent of both. We then selected the best model regarding the effects on occupancy and tested whether transition parameters of poachers depended or not on wildlife presence.

Finally, we considered a second set of hypotheses specific to wildlife response to poachers and environmental covariates. We tested whether wildlife occupancy was conditional to poachers’ occupancy and environmental covariates including ranger patrolling effort, or was only conditional to the presence of poachers, or independent of both (see Appendix A). We did not test spatial or temporal variations in environmental covariates on the transition parameters to respect the closure assumption of the population.

In particular, we expect that under the “wildlife winning” hypothesis, wildlife species are using habitat regardless of patrolling or poaching, indicating that animal space use was more strongly associated with resource distribution than anthropogenic disturbances. Therefore, we expect to find wildlife transition parameters indifferent to poachers’ occupancy and patrolling effort. In contrast, under the “poacher winning” hypothesis, we predict that the presence of poachers will be positively associated with the extinction probability of wildlife and negatively associated with the colonization probability of wildlife.

For each step of the model selection, we used the Akaike information criterion (AIC, Burnham and Anderson 2002) to determine which effects best explain the variation in the data. We considered the model having the lowest AIC to be best supported by the data. We used R software environment (R Core Team 2017) for all analyses. The maximum likelihood estimates were obtained using the quasi-newton method of the optim function in R. We provide the data and R codes in GitHub at https://github.com/oliviergimenez/poaching_occupancy.

From the estimated probabilities of initial occupancy and the estimated transition matrix, we assessed temporal changes in the probability of occupancy in the four states over the study period. We obtained confidence intervals using a parametric bootstrap (Davison and Hinkley 1997). As explained in the introduction we predict an increase in poacher’s occupancy and decrease in wildlife occupancy if the model selection supported the “poacher winning” hypothesis, assuming that we would then use parameter estimates from this model to parametrize those projections.

## Simulations

To validate the performance of our model, we assessed the bias and precision in parameter estimates using two simulation studies focusing on 1) variation in the occupancy design and 2) variation in the sparseness of the data owed to differences in species detectability and occupancy (Appendix C). A previous simulation study on occupancy modelling for a single species revealed that the optimal design for monitoring a rare species was to sample more sites with fewer surveys, and the one for monitoring a common species was to sample few sites with more surveys (MacKenzie and Royle 2005). In our first simulation analysis, we asked the question whether this general recommendation still holds when considering occupancy of two species with different ecology: one with restricted range (wildlife) and the other with a more widespread distribution (poachers) In the second simulation study, we asked which level of sparseness in the data may generate bias in parameter estimation when monitoring species, more or less difficult to detected and with a more or less extended distribution

## Results

### Occupancy data

Over the study period, we collected 322 presence signs of our six key wildlife species (119 of banteng, 119 of barking deer, 9 of gaur, 65 of wild boar, 2 of eld’s deer and 8 of sambar deer) and 377 signs of poachers from the two study areas. We counted 195 detections of poachers only, 213 of wildlife only and 38 detections of both poachers and wildlife in the same sites (Fig.1). Finally, the occupancy table was also composed of 3442 non-detections (Appendix B, Table B1).

### Model selection

The model best supported the “poacher winning” hypothesis and showed the lowest AIC. We found that detection probabilities were related to the spatial and temporal variations of patrolling effort (Table 2) whereas poachers’ occupancy was independent of wildlife distribution and unrelated to rangers’ patrolling effort (AIC = 3105.28, Table 3). We also found that poachers’ transition probabilities were not affected by wildlife, meaning that the dynamics in poacher space use were independent of wildlife distribution. Conversely, we found a positive relationship between extinction probability of wildlife and the presence of poachers and a negative relationship between colonisation probability of wildlife and the presence of poachers (Table 3.1). In the second set of hypotheses testing, focusing on wildlife distribution, the best model showed a positive effect of river length on wildlife occupancy and negative effects of poachers on the dynamics in wildlife space use (Table 3.2, Appendix A). This model was not following “wildlife winning” predictions and had less empirical support compared to the “poacher winning” model from the first set of hypotheses (Table 3).

**Table 2:** Model selection results on detection process. The model names describe the effects of patrolling effort, of roads, rivers and stations, and species interaction (detection is conditional on whether or not species co-occurred) on detection probabilities (see appendix A for corresponding model formulation). The number of parameters, the deviance, AIC and the difference of AICs between the best model and any other candidate models (AAIC) are also indicated.

| Model name | # Parameters | Deviance | AIC | $\Delta AIC$ |
| --- | --- | --- | --- | --- |
| Effects on detection probabilities |  |  |  |  |
| Spatio-temporal effect of patrolling effort on both poachers and wildlife | 21 | 1527.21 | 3112.085 | 0 |
| Distance to road effect on both | 21 | 1554.65 | 3166.97 | 54.88 |
| Constant | 16 | 1563.34 | 3167.189 | 55.11 |
| Spatial effect of patrolling effort on both | 21 | 1563.66 | 3167.824 | 55.74 |
| Species interaction effect | 18 | 1566.38 | 3173.264 | 61.18 |
| Distance to station effect on both | 21 | 1559.45 | 3176.56 | 64.47 |
| Temporal effect of patrolling effort on both | 21 | 1568.71 | 3177.928 | 65.84 |
| Stream length effect on both | 21 | 1561.95 | 3181.57 | 69.48 |

**Table 3:**
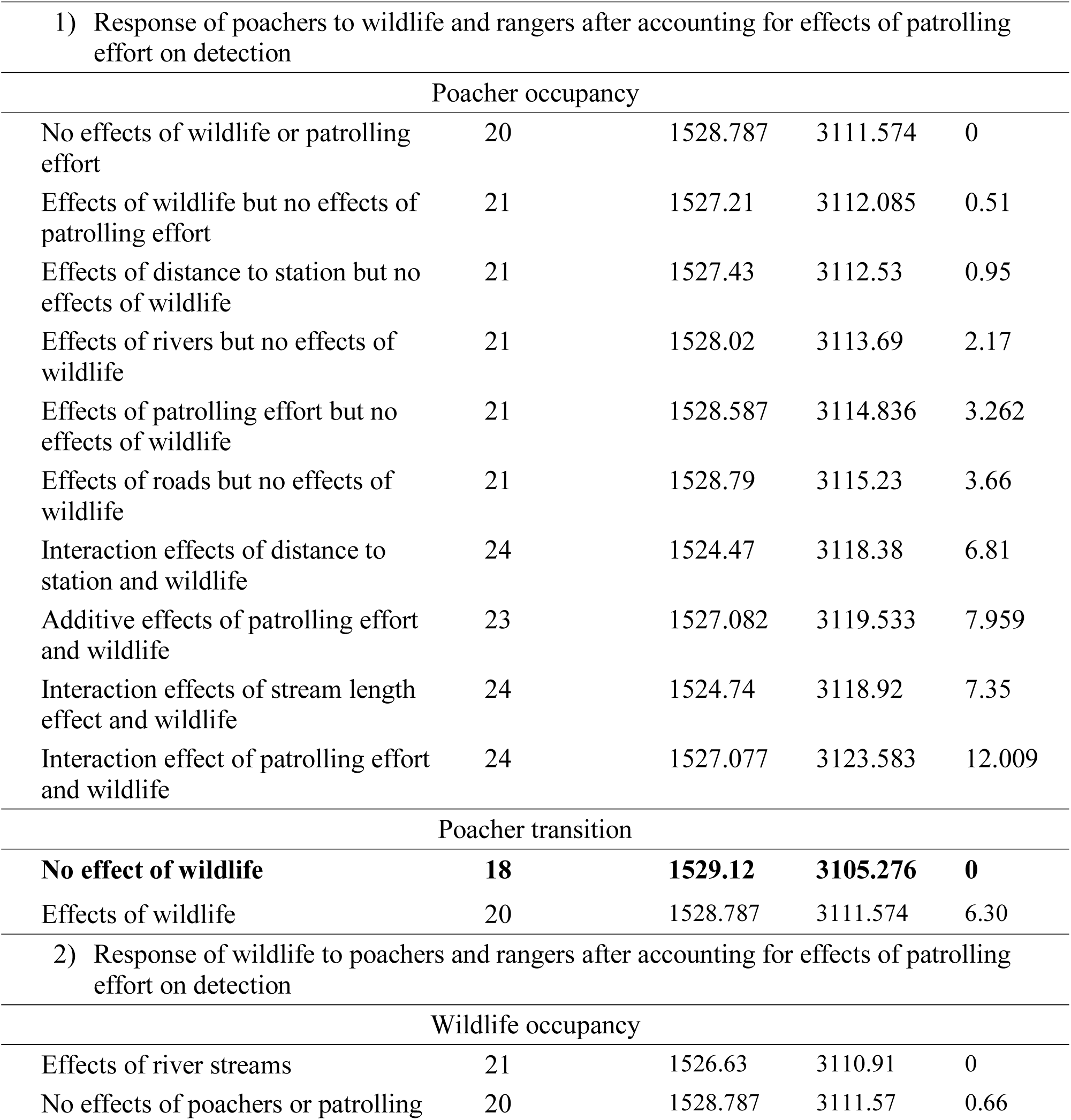
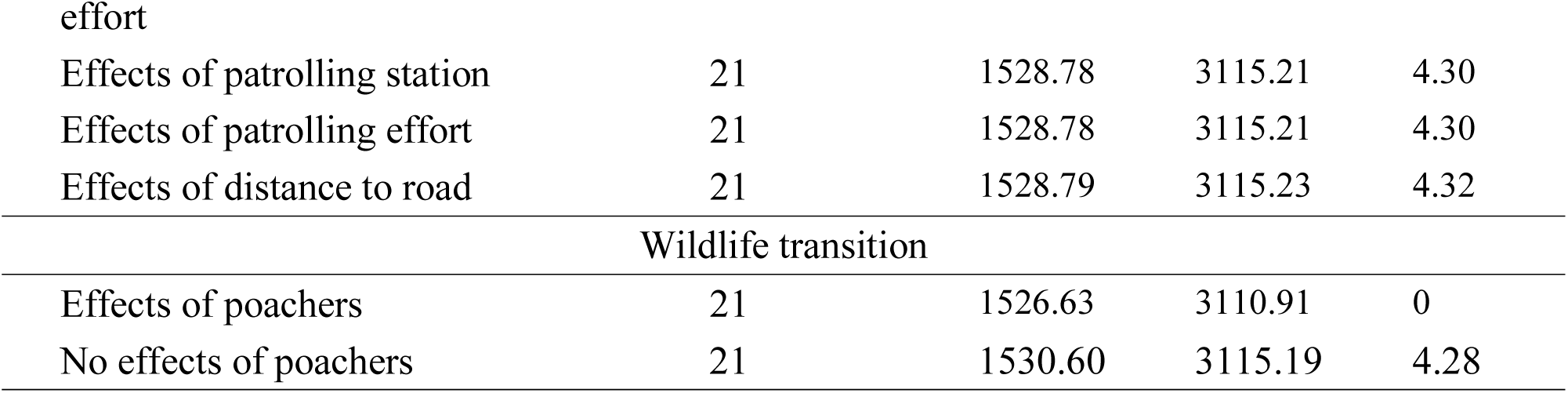
Model selection results on occupancy and transition process using the best model structure for the detection process selected in table 2. Here models for species occupancy and transition are grouped according to our two set of hypotheses: 1) specific to poachers to wildlife occupancy, to rangers (patrolling effort, distance to station), and environmental covariates (roads, rivers), and species interaction (detection is conditional on whether or not species co­occurred), 2) specific to wildlife response to poachers occupancy and environmental covariates (see appendix A for corresponding model formulation). The number of parameters, the deviance, AIC and the difference of AICs between the best model and any other candidate models (AAIC) are also indicated. The best model, highlighted with bold character is the one with the lowest AIC.

The initial occupancy probability of poachers (0.06 ±0.11) was lower than the initial occupancy of wildlife (0.46 ±0.11) (Figure 2). We found that the extinction probability of wildlife in a given year in sites occupied by poachers *(ϵ_w/P_* = 0.80 ± 0.14) was more than double than in sites where poachers were absent *(ϵ_w_/_P_* = 0.36 ± 0.11). The probability of poacher extinction was unrelated to wildlife occurrence *(ϵ_P_/w* = *e_P_/w* = 0.20 ± 0.14). The probability of wildlife colonisation was nearly six times greater in absence of poachers *(γ_w_/p=* 0.28 ± 0.02 with *γ*_wp_ ∼ 0) than in their presence (*γ*_w/p_=0.05 ± 0.02). The probability that a site occupied by wildlife was replaced by poachers (ω_WP_=0.44 ± 0.11) was more than three times the probability that a site occupied by poachers was replaced by wildlife (ω_PW_=0.12 ± 0.11) (Fig. 3). Poachers were also three times more likely than wildlife to remain in a site (site fidelity of 0.63 ± 0.09 for poachers vs 0.20 ±0.11 for wildlife) (Fig. 2). The probability that poachers colonise sites was independent of wildlife present and was very low (*γ*_*P*/*w̄*_ *γ*_*p/w*_ = 0 ±0.11 with *γ*_*WP*_ ∼ 0).

**Figure 2:**
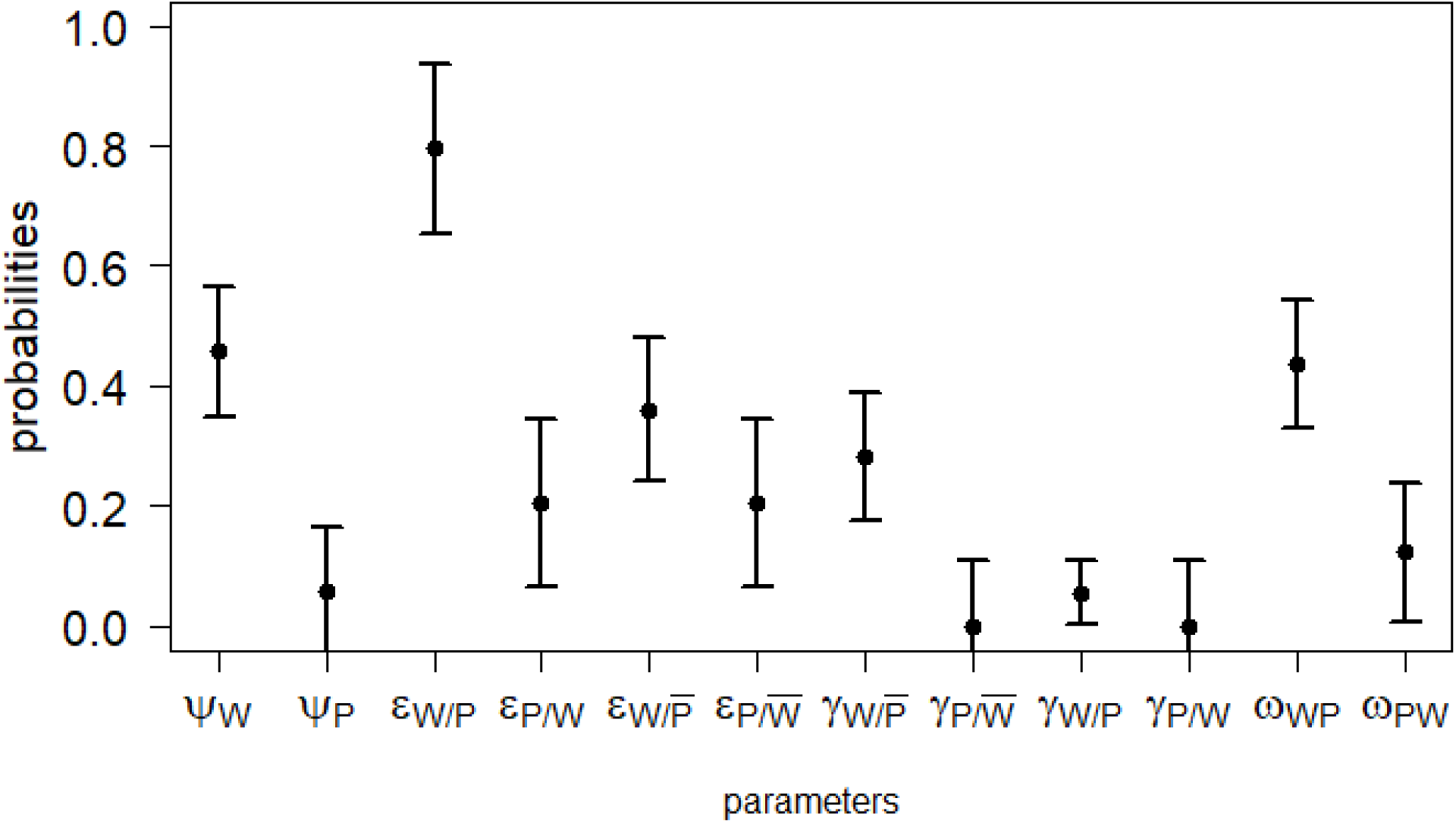
Estimated ecological parameters with associated 95% confidence intervals. We provide the estimated occupancy probability for wildlife (__W_) regardless of the presence/absence of the poachers, occupancy probability for poachers which was independent of wildlife occupancy so we denoted it as (Dp) the extinction probability (ε) of a species (wildlife denoted by W or poachers noted by P) given the presence (without an upper bar) or absence (denoted by an upper bar) of the other (W or P), the colonisation probability of a species (γ) conditional on the presence/absence of the other, the replacement probability of one species by another (ω). Estimates were obtained from the best model displayed in bold character in Table 3 with no effects of wildlife on the transition parameters of the poachers nor on their initial occupancy and spatiotemporal effect of the patrolling efforts on the detection probability of poachers and wildlife.

**Figure 3:**
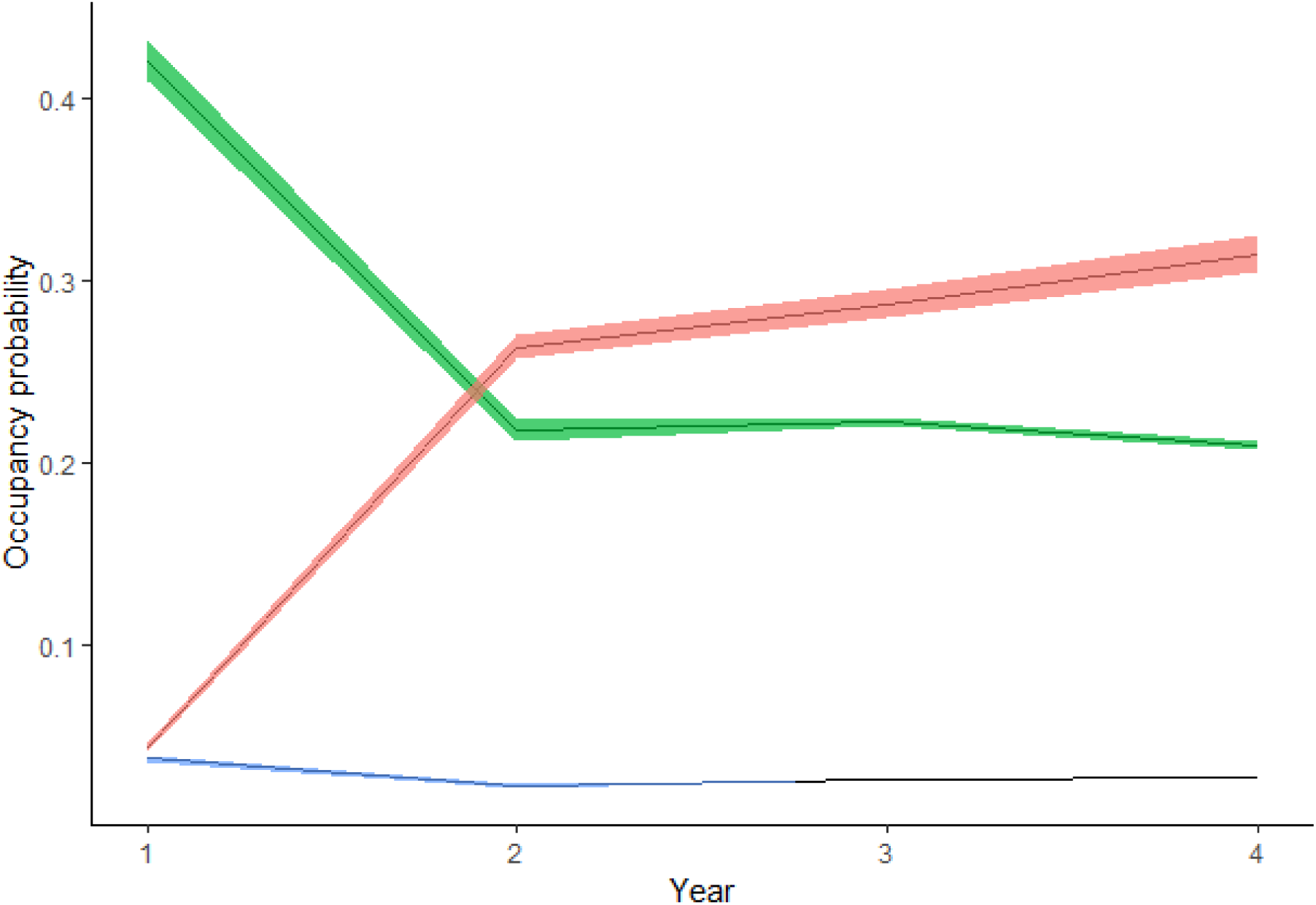
Temporal changes in the probability that a site is occupied by poachers only (OP, in red) and occupied by wildlife only (OW, green), or occupied by both, poacher and wildlife (WP, blue), estimated over the study period.

As predicted by “poachers winning” hypothesis, the estimated probability of sites being occupied by wildlife only was initially high (0.43 +/- 0.02) then decreased by about 50% over the years of the study to 0.22 +/- 0.01. The probability of sites occupied by poachers only exceeded the probability of wildlife occupancy in just the second year of the study. More specifically, it increased from 0.04 +/- 0.01 to 0.30 +/- 0.05 during the study period. The probability of sites occupied by both wildlife and poachers remained relatively constant over the study period (from 0.04 to 0.02 +/- 0.01) (Fig. 3).

Detection probability of poachers and wildlife at the same time was greater than detection of poachers only or wildlife only, and increased rapidly with patrolling effort. The detection probability of wildlife and poachers during a given occasion where both co-occurred was perfect where the patrolling effort was above 0.4. In other words, at sites in which the number of patrol surveys was above 40% the maximum number of GPS waypoints per site were recorded for both poachers and wildlife. The detection probability of *poachers only* increased from 0.04 to 0.96 with increased patrolling effort and was consistently greater than the detection of *wildlife only*, which increased from 0.04 to 0.62 also with increased patrolling effort (Appendix A Fig. A1.1 and A1.2).

## Discussion

The spread of poaching through time and space is an urgent conservation issue; yet its impact on wildlife species remains poorly known even in surveillance systems handled by rangers (Milner-Gulland and Leader-Williams 1992). Here, we combined predator-prey theory of competition for space with a two-species occupancy model to study poacher-wildlife dynamics in protected areas of Cambodia. Our findings largely support the hypothesis that poachers were winning the “space race,” indicated by three lines of evidence. First, from 2013 to 2016, there was a substantial increase in the proportion of sites occupied only by poachers (approximately from 80 to 600 km^2^ in PPWS and from 148 to 1110 km^2^ in SWS as explained in Appendix A). Second, over the same time period, there was a dramatic decline of the spatial coverage by wildlife. Third, the reduction in wildlife occupancy between years was directly attributable to the presence of poachers, as indicated by both the “extinction” and “colonisation” parameters. If poaching was reduced by half, for example, our models indicate that the occupancy range of the wildlife detected in our study could gradually increase from 400 km^2^ to 580 km^2^ in PPWS and from 740km^2^ to 1073 km^2^ in SWS (Appendix A). Combined, these lines of evidence suggest that the increasing presence of poachers in our two study sites has substantially reduced the spatial occupancy of ungulate species, with serious consequences on prey availability for tigers.

Although we found support for the “poachers winning” hypothesis, some of our results do not directly align with theories of wild predator-prey space use. Previous research on predator-prey space use in a landscape of fear established that greater levels of spatial overlap between predators and prey indicate that predators are winning the race. Whereas, lower levels of spatial overlap indicate that prey are winning the race (Sih 2005). However, in our case study, we found that the proportion of sites occupied both by poachers and wildlife (“WP” in Fig. 3) remained relatively low and constant. This difference might be due to poachers operating very differently from wild predators. Crucially, unlike wild predators, many poachers in this area are hunting for economic gain not individual subsistence (Louck et al. 2009). Because poachers seek to maximize their offtake relative to effort, for example, through blanket use of methods that capture and kill a wide-range of species, the per-capita impact of poachers on wild ungulates is likely higher than that of wild predators. As a result, wild predators might be more likely to co­occur with wild prey at the site-level, whereas the relatively fast and intense poaching pressure we observed in our study completely suppressed wild ungulate occupancy.

Another factor that differentiates our study from a typical study on wild predator-prey systems is the presence of rangers. For example, rangers change the distribution of snares (i.e., predation pressure) through removal and could alter the behaviors of poachers and wildlife due to their patrolling routes and schedules. In addition, wildlife could use patrolling stations or roads as “shields” to avoid poachers, similar to prey species sometimes using areas close to human activities (e.g., settlements) as refuge from wild predators (Muhly et al. 2010). However, in our case study, wildlife species were probably unable to differentiate between poachers and rangers, and we did not find evidence for rescue effects arising from ranger patrols. Instead, the positive effect of river density on wildlife occupancy suggests wildlife were selecting for areas with high quality habitat (Dudgeon et al. 2000), despite the top-down pressure from poachers. The extent to which predator-prey theories advance our understanding of poacher-wildlife systems is a fruitful area of future research. Insights from such work can generate testable hypotheses that ultimately improve the effectiveness of anti-poaching interventions.

Our results also suggest that the patrolling effort in the PPWS and SWS study areas of Cambodia are currently inadequate, in both the spatial coverage and the time allocation of rangers’ forces. Increasing patrolling effort improved detection but not occupancy probabilities of poachers, meaning that increasing the frequency of survey within a site was effective for detecting snares but not for preventing poaching. Poachers tend to put more snares in sites where wildlife is abundant in order to maximize their probability of capture success. Doing so likely also increases the chance of their snares being detected and removed by rangers. This could explain why the detection probability of both poachers and wildlife was greater than the detection probability of poachers only or wildlife only (see Hines et al. 2010 for effects of abundance on detection). Our results therefore suggest that there is a diminishing return on investment beyond a certain patrolling effort. Instead of exceeding that effort level, ranger resources would be better invested in monitoring new sites. In particular, increased patrolling of those sites where only poaching-related threats occur may stem the invasion of poachers to new sites. It is important to note, however, that without effective criminal prosecution of poachers, those individuals might continue to poach despite the detection of their snares and activities.

### Model limitations

The strength of some of our inferences is limited because, unfortunately, we could not estimate the occupancy of wildlife only despite the presence of poachers and vice versa (see Fidino et al 2018, Miller et al. 2012). We ran preliminary analysis using the same model as in Miller et al. 2012, where detection probability of one species was conditional on the presence of the other. Unfortunately, our data was insufficient and too sparse to estimate such parameters. Therefore, we opted for a simpler representation, prioritizing the tests on the effects of patrolling effort on the detection of both wildlife and poachers. Also, we acknowledged that pooling six species of ungulates into one guild defined as “tiger prey” could also generate bias in estimates, as each species has its own abundance, occupancy dynamics and detection probabilities. To address this problem in the future, statistical ecologists could develop a mixture dynamic occupancy model within the multi-species framework to deal with heterogeneity in detection and occupancy (MacKenzie et al. 2018)

Our model relied on several assumptions, which under certain circumstances, can be violated. We assumed the population to be closed during the 12-months’ primary occasion. However, poaching-related threats may change within a year given the incentive of poachers to displace snares after catching an animal and given the role of rangers to remove snares every time they detect one. This could overestimate the occupancy of poachers in a single-species model (Rota et al. 2009). This is the reason why we viewed “site-occupancy” as “site-use” by a species, similar to Moore et al. (2017).

We did not account for spatial autocorrelation among detections and occupancy states, which is likely to occur in interacting species, monitored with transects (Hines et al., 2010, Guillera-Arroita et al., 2011). Our model also assumes spatial independence in the process governing the dynamics across the species’ entire range. This assumption can be relaxed via Bayesian methods using estimated occupancy of neighbouring sites in the previous time step as a covariate to predict extinction/colonisation processes in the current step (Green et al., 2018; Heard et al., 2013) or mixture dynamic occupancy models as mentioned above.

Finally, assessing the quality of fit of occupancy models accounting for imperfect detection is not straightforward. In standard species distribution models, AUC metrics are routinely used in routine for model evaluation and predictive performance. AUC metrics can still be computed in occupancy models under imperfect detection, but they no longer assess how well it predicts true occupancy but rather how well it predicts detection (Lahoz-Monfort et al. 2014). MacKenzie and Bailey (2004) developed a goodness-of-fit test for single-season occupancy models for one species only but no test has so far been developed for two-species occupancy models. The simulations we performed in a two-fold validation process, demonstrated that our model was appropriate for the sampling design and the community dynamics in our case study,, but we encourage future research into goodness-of-fit tests for the models we have used.

### Conclusions

To our knowledge, this study is one of the few applications of a two-species occupancy model extended to a multi-season version (see also Fidino et al., 2018, Yackulick et al. 2014). It is also the first to apply such a model to explicate the space use dynamics of a poacher-wildlife system. As such, our model helps quantify the rate at which poaching pressure shrinks the occupied range of wildlife species, as well as wildlife communities, in relatively data-poor systems. Range contraction is an important indicator of species extinction risk (Burgess et al. 2017). Our study is also a proof of concept for a science-based conservation program in the wildlife sanctuaries of Cambodia. We can use our results to parameterize a predictive distribution model and implement structured-decision making (SDM) to determine cost effective patrolling strategies for rangers in Cambodia (see Martin et al. 2001 combining occupancy modelling and SDM). Such a program is needed, for example, to help understand how poaching-induced prey depletion will influence tiger reintroduction efforts in the region. Furthermore, because our model utilized SMART data from ranger patrols, it can be easily replicated in many regions around the world that use that use the SMART system. Continued development of methods for evaluating wildlife distribution dynamics that account for the impacts of illegal hunting, as with our model, will enable better prediction of how different poaching interdiction strategies influence wildlife recovery in space and time.

## Acknowledgments

We are grateful to SMART partnership, WWF Cambodia, the ministry of Environment Cambodia, along with the rangers who have planned and collected the data from the Phnom Prich wildlife sanctuary and Serepok Sanctuary and the collaboration of the Asia Poaching prevention group of WWF. LM and OG were funded by the French National Research Agency (grant ANR-16-CE02-0007). We thank the reviewers for their helpful and constructive suggestions and proposed corrections to improve our study.

