## supplemental material for "Inferring wildlife poaching in Southeast Asia with multispecies dynamic occupancy models"

**Appendix A**

**Supplementary information on the model structure and data analyses**

Our two-species dynamic occupancy model allowed assessing the direction and intensity of the interactions, disentangled from other environmental factors. First we considered the Richmond-Waddle parameterization on occupancy and detection to better assess the contribution of environmental and anthropogenic covariate on the detectability of poachers and wildlife and the spatial overlap between both. Indeed this parameterization is numerically more stable and more appropriate when covariates are included as the regression coefficients for these covariates don’t need to be formulated in a generalized logit scale (Yackulic et al. 2014, MacKenzie et al. 2017). Second, we quantify two-way spatial interactions by estimating extinction/colonization probabilities of a species guild (wildlife or poacher) conditional to the occupancy of the other.on a yearly dynamic, which few studies have done so far (Cusack et al. 2018).

First, we accounted for the different effects on the detection process. We provided the structure of each competing model where the probability of detection can be expressed in a generalized linear modelling spirit allowing the probability of interest to be related to environmental covariates (patrolling effort X, distance to road DR, distance to station DS and length of river stream LS), using a logit link function. We considered the following models using the same name as in Table 1 in the main text:

- *Constant model*: poachers and wildlife species are similarly detectable so they have the same detection probabilities

$logit\left( p_{WP} \right)=logit\left( p_{W} \right)=logit\left( p_{P} \right)$ = β_0_

$$or$$

$$p_{WP}=p_{W}=p_{P}=\beta_{0}$$

- *Species interactions effects*: detection is conditional on whether the species co-occurred or not with the other

$$p_{W}=\mathrm{logit}^{-1}(\beta_{1})$$

$$p_{P}=\mathrm{logit}^{-1}(\beta_{2})$$

$$p_{WP}=\mathrm{logit}^{-1}(\beta_{3})$$

- *Spatio-temporal effect of patrolling effort* on the detection probability of poachers only, wildlife only, and of both poachers and wildlife: Spatial-temporal variation in detection as a function of the monthly patrolling effort per site. In other words, we estimate a detection probability for each site *i* at time *t* (with t a sampling occasion at a primary occasion k and secondary occasion j)

$$p_{W}(i,t)=\mathrm{logit}^{-1}(\beta_{1} + \beta_{2}X_{i,t})$$

$$p_{P}(i,t)=\mathrm{logit}^{-1}( \beta_{3}+ \beta_{4}X_{i,t})$$

$p_{WP}(i,t)=\mathrm{logit}^{-1}( \beta_{5}{.Z}_{i,k}^{WP}+ \beta_{6}.X_{i,t}$)

with $X_{i,t}$the frequency of rangers visits per site i and sampling occasion t as a measure of patrolling effort

- *Temporal effect of patrolling effort* on the detection probability of poachers only, wildlife only, and of both poachers and wildlife: detection is a function of monthly variations in patrolling effort.

$p_{W}(t)=\mathrm{logit}^{-1}(\beta_{1} + \beta2.X_{.,t})$

$$p_{P}(t)=\mathrm{logit}^{-1}(\beta_{3}+\beta_{4}X_{.,t})$$

$$p_{WP}(t)=\mathrm{logit}^{-1}(\beta_{5} + \beta_{6}X_{.,t})$$

with $X_{.,t}$the mean frequency of rangers visits across sampling occasions during sampling occasion *t*

- *Spatial effect of patrolling effort* on the detection probability of poachers only, wildlife only, and of both poachers and wildlife: detection depends on the patrolling effort across sites. Rangers patrolling site *i* has a specific chance to detect a species given that the species occupies the site.

$$p_{W}(i)=\mathrm{logit}^{-1}(\beta_{1} + \beta_{2} X_{i,.})$$

$$p_{P}(i)=\mathrm{logit}^{-1}(\beta_{3} + \beta_{4} X_{i,.})$$

$$p_{WP}(i)=\mathrm{logit}^{-1}(\beta_{5} + \beta_{6} X_{i,.})$$

with $X_{i,.}$the average frequency of rangers visits per site i across sampling occasions

- *Distance to road effect* on the detection probability of poachers only, wildlife only, and of both poachers and wildlife: spatial variation in detection of poachers only, wildlife only, and both were explained by distance to road

$$\mathrm{logit}\left( p_{W,i} \right)=\mathrm{logit}^{-1}(\beta_{1} + \beta_{2} \mathrm{DR}_{i})$$

$$logit(p_{P,i})={\mathrm{logit}^{-1}(\beta}_{3} + \beta_{4} \mathrm{DR}_{i})$$

$$logit(p_{WP,i})={\mathrm{logit}^{-1}(\beta}_{5} + \beta_{6} \mathrm{DR}_{i})$$

with DRi the minimum distance between the center of site i and the nearest road

- *Distance to station effect* on the detection probability of poachers only, wildlife only, and of both poachers and wildlife in a given site *i* is a linear function of the closest distance to ranger station.

$$p_{W}={\mathrm{logit}^{-1}(\beta}_{1} + \beta_{2}\mathrm{DS}_{i})$$

$$p_{P}={\mathrm{logit}^{-1}(\beta}_{3}+ \beta_{4}\mathrm{DS}_{i})$$

$$p_{WP}=\mathrm{logit}^{-1}(\beta_{5}+ \beta_{6}\mathrm{DS}_{i})$$

with DSi the distance between the center of each site *i* and the closest patrolling station

- *Stream length effect* on the detection probability of poachers only, wildlife only, and of both poachers and wildlife in a given site *i*:

$$p_{W,i}={\mathrm{logit}^{-1}(\beta}_{1} + \beta_{2} \mathrm{SL}_{i})$$

$$p_{P,i}=\mathrm{logit}^{-1}(\beta_{3} + \beta_{4} \mathrm{SL}_{i})$$

$$p_{WP,i}={\mathrm{logit}^{-1}(\beta}_{5} + \beta_{6} \mathrm{SL}_{i})$$

with *SL*_i_ the length of stream crossing site *i*

We used the best model structure for the detection process selected in the preceding step (*Spatio-temporal effect of patrolling effort on both* with six regression coefficients associated to detection probabilities) and tested two sets of biological hypotheses regarding: 1) poachers response to wildlife, rangers and environmental covariates 2) response of wildlife to poachers and rangers and environmental covariates.

1) We first focused on poachers’ occupancy considering the effects of wildlife presence/absence, rangers patrolling effort and other factors.

- *No effects of wildlife or patrolling effort*: occupancy of poachers regardless of the presence or absence of wildlife

$$\psi_{P/W}={\mathrm{logit}^{-1}(\beta}_{7})$$

$$\psi_{P/\bar{W}}=\mathrm{logit}^{-1}(\beta_{7})$$

$$\psi_{W}={\mathrm{logit}^{-1}(\beta}_{8})$$

- *Effects of wildlife no effects of patrolling effort* here we assume the occupancy probability of poachers is conditional on presence/absence of wildlife

$$\psi_{P/W}={\mathrm{logit}^{-1}(\beta}_{7})$$

$$\psi_{P/\bar{W}}={\mathrm{logit}^{-1}(\beta}_{8})$$

$$\psi_{W}={\mathrm{logit}^{-1}(\beta}_{9})$$

- *Effects of patrolling effort but no effects of wildlife*: spatial effect of patrolling effort on poacher occupancy only

$\psi_{P/W}={\mathrm{logit}^{-1}(\beta}_{7}+ \beta_{8} X_{i,t})$

$$\psi_{P/\bar{W}}={\mathrm{logit}^{-1}(\beta}_{7}+ \beta_{8} X_{i,t})$$

$$\psi_{W}=\mathrm{logit}^{-1}(\beta_{9})$$

- *Effects of distance to station but no effects of wildlife* on poachers’ occupancy probability

$\psi_{P/W}=\mathrm{logit}^{-1}(\beta_{7} + \beta_{8}\mathrm{DS}_{i})$

$$\psi_{P/\bar{W}}={\mathrm{logit}^{-1}(\beta}_{7} +\beta_{8} \mathrm{DS}_{i})$$

$$\psi_{P/\bar{W}}=\mathrm{logit}^{-1}(\beta_{9})$$

- *Effects of distance to road but no effects of wildlife* on poachers’ occupancy probability

$\psi_{P/W}={\mathrm{logit}^{-1}(\beta}_{7}+ \beta_{8} \mathrm{DR}_{i})$

$$\psi_{P/\bar{W}}={\mathrm{logit}^{-1}(\beta}_{7}+ \beta_{8} \mathrm{DR}_{i})$$

$$\psi_{W}=\mathrm{logit}^{-1}(\beta_{9})$$

- *Effects of stream length but no effects of wildlife* on poachers’ occupancy probability

$\psi_{P/W}={\mathrm{logit}^{-1}(\beta}_{7}+\beta_{8}\mathrm{LS}_{i}$ )

$$\psi_{P/\bar{W}}=\mathrm{logit}^{-1}(\beta_{7} + \beta_{8} \mathrm{LS}_{i})$$

$$\psi_{W}=\mathrm{logit}^{-1}(\beta_{9})$$

- *Interaction effects of patrolling effort and wildlife*: poacher occupancy conditional in presence of wildlife varies with patrolling effort across sites and poacher occupancy conditional to the absence of wildlife

$\psi_{P/W}={\mathrm{logit}^{-1}(\beta}_{7}+\beta_{8}X_{i,t})$

$$\psi_{P/\bar{W}}={\mathrm{logit}^{-1}(\beta}_{9}+\beta_{10}. X_{i,t})$$

$$\psi_{W}={\mathrm{logit}^{-1}(\beta}_{10})$$

- *Interaction effects of distance to station and wildlife*

$\psi_{P/W}={\mathrm{logit}^{-1}(\beta}_{7}+ \beta_{8}. \mathrm{DS}_{i})$

$$\psi_{P/\bar{W}}=\mathrm{logit}^{-1}(\beta_{9} + \beta_{10}. \mathrm{DS}_{i})$$

$$\psi_{W}={\mathrm{logit}^{-1}(\beta}_{11})$$

- *Interaction effects of distance to road and wildlife*

$\psi_{P/W}={\mathrm{logit}^{-1}(\beta}_{7}+ \beta_{8}\mathrm{DR}_{i})$

$$\psi_{P/\bar{W}}={\mathrm{logit}^{-1}(\beta}_{9}+ \beta_{10} \mathrm{DR}_{i})$$

$$\psi_{W}={\mathrm{logit}^{-1}(\beta}_{11})$$

- *Interaction effects of stream length and wildlife*

$\psi_{P/W}=\mathrm{logit}^{-1}(\beta_{7}+ \beta_{8}. \mathrm{LS}_{i})$

$$\psi_{P/\bar{W}}={\mathrm{logit}^{-1}(\beta}_{9}+ \beta_{10}. \mathrm{LS}_{i})$$

$$\psi_{W}={\mathrm{logit}^{-1}(\beta}_{11})$$

Then we kept the best structure on occupancy of poachers and tested effects on transition parameters (See the script “transition_effects” on the github repository for more details on model structure). Here, the transition probabilities follow a generalized multinomial logit function where each parameter describing a transition between two states of site occupancy is function of the regression coefficients specific to the other states occupancy. In other words, the sum between columns of each row of the transition matrix T must equal to 1. In contrast to Miller et al. 2011 or Yackulic et al. 2014, our parameterization describes the transition from unoccupied site (U) to occupied by both (WP) by using only one parameter, $\gamma_{WP}$, and transition WP to U, $\varepsilon_{WP}$, instead of the product of the two species colonizing separately, $\gamma_{W}\gamma_{P}$ or $\varepsilon_{P} \varepsilon_{W}$. Indeed, we found a stronger interest in accounting for one parameter per matrix element. This way we could easily test hypotheses on the poacher-wildlife space race under patrolling effort. This was done by looking at the underlying mechanism in the transition probability of poachers in presence or absence of wildlife and transition probability of wildlife given the presence or absence of poachers, rather than partitioning the process when both are simultaneously going extinct or colonizing which is less likely to occur under ranger patrol. Indeed, when both wildlife and poachers are present we expected rangers to either succeed in extirpating poachers and saving wildlife or failing to protect wildlife, which may go extinct locally while poachers remain.

We started by looking at effects of wilflide presence on poachers transition probabililities

*Effects of wildlife* on poachers’ transition parameters

$\gamma_{W/\bar{P}}$ = e^β13^ / (1+ e^β12^ + e^β13^ + e^β14^ )

$\gamma_{P/\bar{W}}$ = e^β12^ / (1+ e^β12^ + e^β13^ + e^β14^ )

$\gamma_{WP}$ = e^β14^ / (1+ e^β12^ + e^β13^ + e^β14^ )

$\epsilon_{WP}$ = e^β9^ / (1+ e^β9^ + e^β10^ + e^β11^ )

$\epsilon_{W/P}$ = e^β10^ / (1+ e^β9^ + e^β10^ + e^β11^ )

$\epsilon_{P/W}$= e^β11^ / (1+ e^β9^ + e^β10^ + e^β11^ )

$\epsilon_{P/\bar{W}}$ = e^β15^ / (1+ e^β15^ + e^β16^ + e^β17^ )

$\omega_{PW}$= e^β16^ / (1+ e^β15^ + e^β16^ + e^β17^ )

$\gamma_{P/W}$ = e^β17^ / (1+ e^β15^ + e^β16^ + e^β17^ )

$\epsilon_{W/\bar{P}}$ = e^β18^ / (1+ e^β18^ + e^β19^ + e^β20^ )

$\omega_{WP}$= e^β19^ / (1+ e^β18^ + e^β19^ + e^β20^ )

$\gamma_{W/P}$ = e^β20^ / (1+ e^β18^ + e^β19^ + e^β20^ )

Under the *“Poachers winner” hypothesis,* we assume poachers’ colonization and extinction independent of the prey occupancy:

$\gamma_{WP}$ = = e^β14^ / (1+ e^β12^ + e^β13^ + e^β14^ )

$\gamma_{P/\bar{W}}$ = = e^β12^ / (1+ e^β12^ + e^β13^ + e^β14^ )

$\gamma_{P/W}$= $\gamma_{P/\bar{W}}$

$\epsilon_{P/W}$ = e^β11^ / (1+ e^β9^ + e^β10^ + e^β11^ )

$\epsilon_{P/\bar{W}}$= $\epsilon_{P/W}$

and we assume wildlife colonization and extinction conditional on presence of poachers

$\epsilon_{W/P}$ = e^β10^ / (1+ e^β9^ + e^β10^ + e^β11^ )

$\epsilon_{WP}$= e^β9^ / (1+ e^β9^ + e^β10^ + e^β11^ )

$\gamma_{W/\bar{P}}$ = = e^β13^ / (1+ e^β12^ + e^β13^ + e^β14^ )

$\gamma_{W/P}$ = = e^β18^ / (1+ e^β11^ + e^β17^ + e^β18^ )

$\epsilon_{W/\bar{P}}$= e^β11^ / (1+ e^β11^ + e^β17^ + e^β18^ )

$\omega_{WP}=$ e^β17^ / (1+ e^β11^ + e^β17^ + e^β18^ )

$\omega_{PW}$ = e^β16^ / (1+ e^β16^ + e^β11^ + e^β12^ )

2) We then proceed similarly for testing the effects of poachers, rangers patrolling effort and other factors on wildlife ecological parameters (see the “occupancy_effects” and “transition_effects” scripts in Github repository).

**Supplementary information on the results**

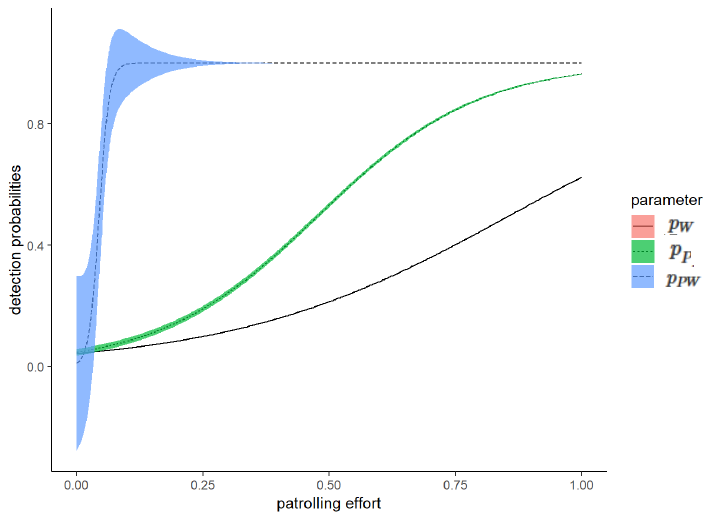

Figure A.1: Estimated site-specific detection probabilities and their 95% confidence intervals as a function of patrolling effort. The figure represents detection of both wildlife and poachers *p_WP_* (dash line line blue ribbon), detection probability of poacher only *p_P_* (dotted line green ribbon), detection probability of wildlife only (solid line). Estimates were obtained from the model best supported by the data, see Table 1 in main text.

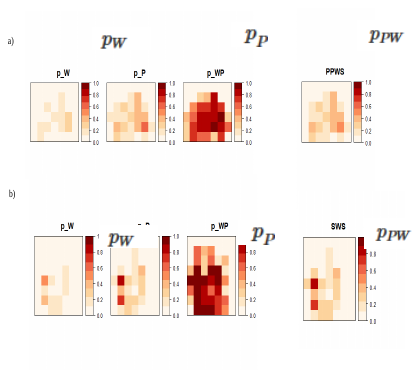

Fig. A.2: Spatial variation in the probability of detection of wildlife only *p_W_*, poachers only *p_P_*, and the detection probability of both wildlife and poachers *p_WP_* in a) Phnom Prich Wildlife Sanctuary (PPWS) b) Srepock Wildlife sanctuary (SWS)

Calculation of the increase in area occupied by poachers and the decrease in area occupied by wildlife:

The initial probability of a site occupied by poachers regardless of wildlife was 0.04. Therefore, poachers initially occupied 4% of the 2000 km^2^ of the Phnom Prich Wildlife Sanctuary that is 80 km^2^. Similarly, poachers initially occupied 4 % of the 3700 km^2^ of the Serepok Wildlife Sancturary that is 148 km^2^. After 4 years, we estimated that sites occupied only by poachers increase from 4 % to 30%. So the area occupied by poachers increase from 80 km^2^ to 600 km^2^ (0.30*2000) in PPWS and from 148 to 1110 (0.3*1110) in SWS. The probability of a site occupied by wildlife decreased from 0.43 to 0.20. Therefore, out of the 2000 km^2^ area in PPWS, wildlife initially occupied 860 km^2^ in 2013 and its distribution shrinked to 400 km2 in 2016. Out of the 3700 km^2^ area in SWS, wildlife initially occupied 1591 km^2^ in 2013 but only 740 km2 in SWS. Now, we assumed poaching reduced by half. In other words, we assumed the site-specific extinction probability of wildlife in presence of poacher $\epsilon_{W/P}$decreased from 0.8 to 0.4. We used Monte Carlo simulations and project distribution ranges of sympatric species for one more year, with the updated value of wildlife extinction probability in the transition matrix. Then, we predicted that 29% of the study areas would be covered by wildlife only, which would correspond to 580km^2^ of the PPWS and 1073 km^2^ of the SWS.

**Appendix B**

**Supplementary information on the data, sampling design and patrolling protocol**

*Data*

We checked the sample size of our occupancy table by first counting the total numbers of observations in the four occupancy states (U, OP, OW, WP). We also determined the number of transitions observed between states during the 4 consecutive years of the study period and reported them in Table 1. We looked at whether both species were detected during a primary occasion (year), meaning that if we found at least OW and OP or WP only during the 12 monthly repeated surveys of a primary occasion we assigned the occupancy state as both species co-occurring. If only OW or OP were reported within a year then we assigned the observation for that primary session as occupied only by wildlife of poachers. If ranger did not detect any information on either wildlife or poachers within a site and during a full year, we consider the site as unoccupied. We data description allowed us to a priori assess whether the number of transition events was sufficient to fit a dynamic two-species occupancy model with this study design.

Here we point out that we found a 0 probability of extinction of both poachers and wildlife in a site occupied by both on a given year, as well as a 0 colonisation probability of both poachers and wildlife. We considered them not estimable, as we recorded only one observation for these transitions associated to those probabilities (one from U to WP, one from WP to U) (Table B.1).

Table B.1 : Number of site occupancy states observed across the 4 years of study. We provide the yearly sample size of the number of observations of sites occupied by wildlife only OW, poachers only OP, both WP and none U and the number of observed transitions from one state to another counted in the occupancy table.

| Events in occupancy states | Total # of events | Sample size per year (within primary occasions) | | | | Observed transition between primary occasions | | | |
| --- | --- | --- | --- | --- | --- | --- | --- | --- | --- |
|  |  | 2013 | 2014 | 2015 | 2016 | OW | OP | WP | U |
| OW | 213 | 70 | 56 | 68 | 19 | 27 | 19 | 13 | 22 |
| OP | 195 | 16 | 35 | 71 | 73 | 4 | 10 | 3 | 8 |
| WP | 38 | 7 | 5 | 14 | 12 | 11 | 4 | 4 | 1 |
| U | 3442 | 879 | 876 | 819 | 868 | 14 | 13 | 1 | 89 |

*Covariates*

All covariates (patrolling effort, stream length crossing each site and minimum distance between the centroid of each site and the nearest road or nearest station) were correlated to some extent. For this reason, we tested the effect of each covariate on parameter estimates in separate models. The two covariates measuring the closest distance to the road and the closest distance to the station were particularly highly correlated, with r^2^ = 0.89 in PPWS and r^2^ = 0.61 in PWS. Otherwise, distance to road was negatively correlated to stream length, with r^2^ = - 0.43 in both study areas. The least correlated covariates were the distance to station and stream length across SWS sites (r^2^ = -0.17).

1. Spatial variation in the patrolling effort in terms of the mean frequency of visits during the study period

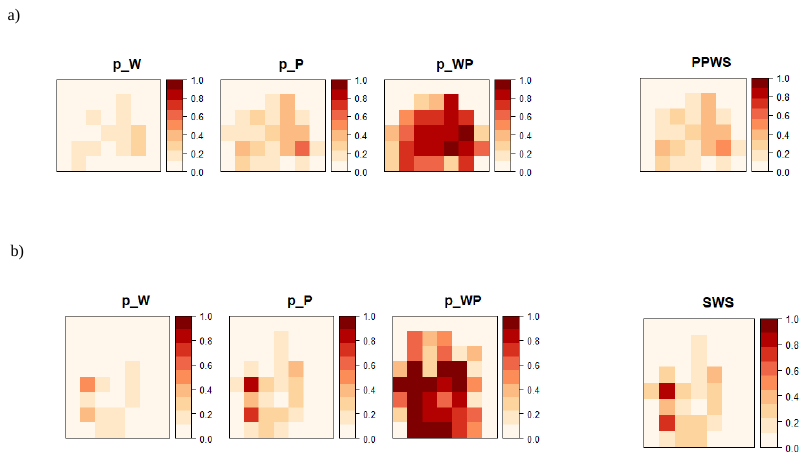

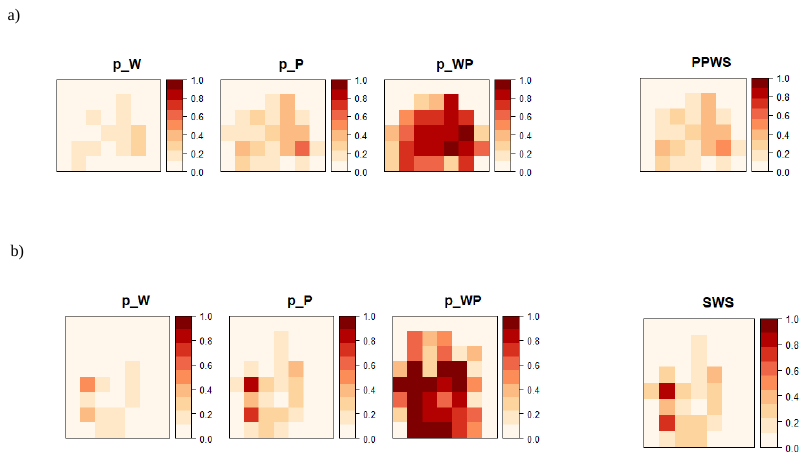

Fig. B.1: Spatial representation of patrolling effort, showing the mean frequency of visits during the study period in the Phnom Prich wildlife sanctuary (PPWS) and the Serepok wildlife sanctuary (SWS).

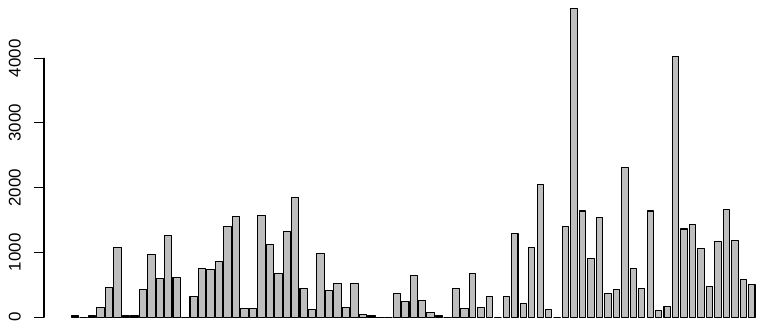

Figure 2.B: histogram of the total number of way points recorded per site during the study period in PPWS.

2. Temporal characteristics of patrolling sessions

Cambodian weather is characterized by two season: monsoon from December until April followed by a dry season. We did not find any inter or intra annual patterns to explain patrolling effort. Variation in patrolling effort was more related to the availability of financial resources. In the PPWS, the total number of waypoints recorded per sampling occasion (per months) varied from 750 in April 2014 to more than 1800 in December (Figure 3.B). The number of daily patrolling sessions within a month range from 0 day to 22 days (Figure 4.B) and the number of waypoints recorded per day across all sites could vary from 2 to 201. All sites were prospected at least once per year. The maximum waypoints recorded within a year per site was 1621, the maximum number of waypoints per month per site was 224 (Figure 5.B).

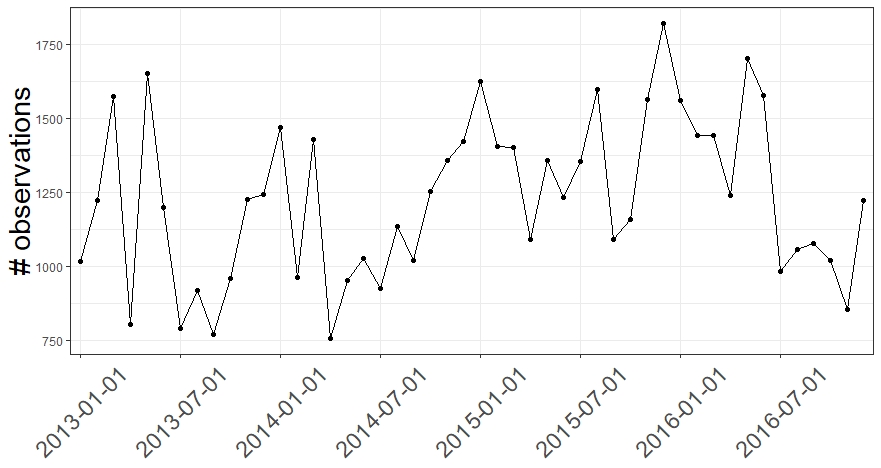

Figure 3.B: number of recorded waypoints per sampling occasions during the 4 years study periods in Phnom Prich Wildlife Sancturary

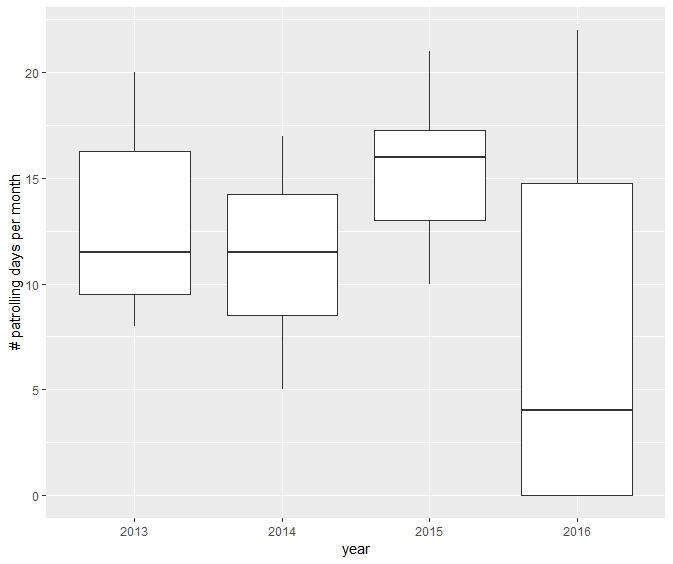

Figure 4.B: Number of daily patrolling sessions per month over the 4 years of the study period. The boxes encompass the first to the third quantiles, the horizontal lines inside the box stand for the median and the whiskers are the interquartile range.

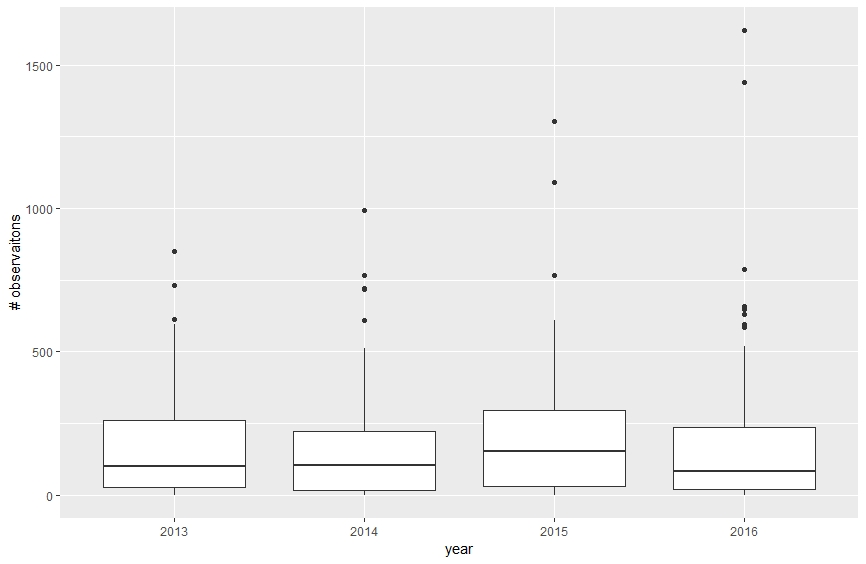

Figure 5.B: Number of GPS waypoints recorded per sites within primary occasions (from 2013 until 2016). Empty circles represent outliers, the boxes encompass the first to the third quantiles, the horizontal lines inside the box stand for the median and the whiskers are the interquartile range.

3. Spatial and temporal variation of GPS waypoints recorded by rangers across the Phnom Prich Wildlife Sanctuary (PPWS) and Serepock Wildlife Sancturary (SWP) during the 4 years of study period

Here after we provide a monthly map of GPS waypoints, when rangers recorded presence signs of animal (in green), presence signs of poaching (in red), nothing detected (in black) per sampling occasion (month) in the PPWS study area.

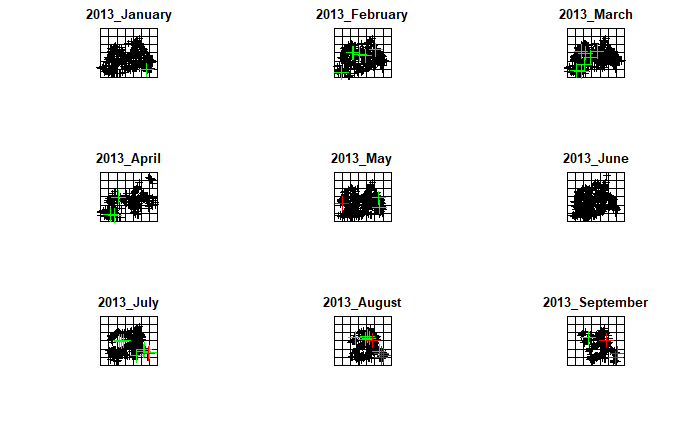

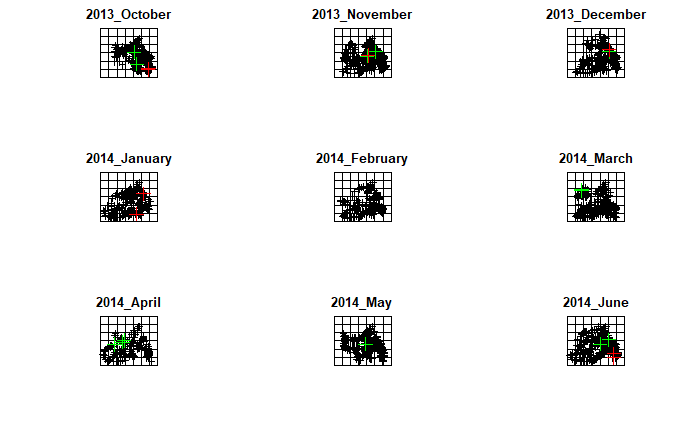

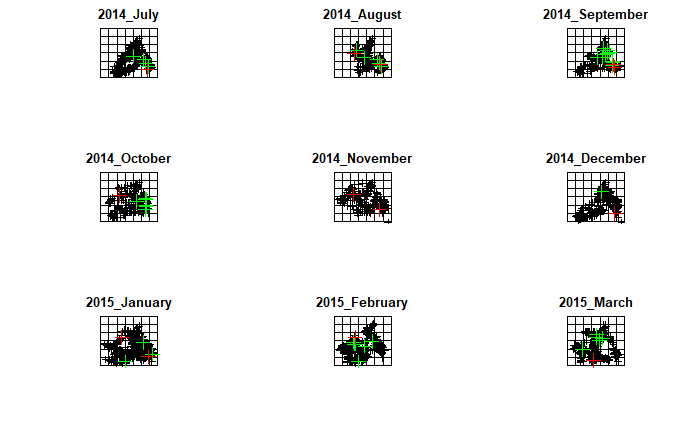

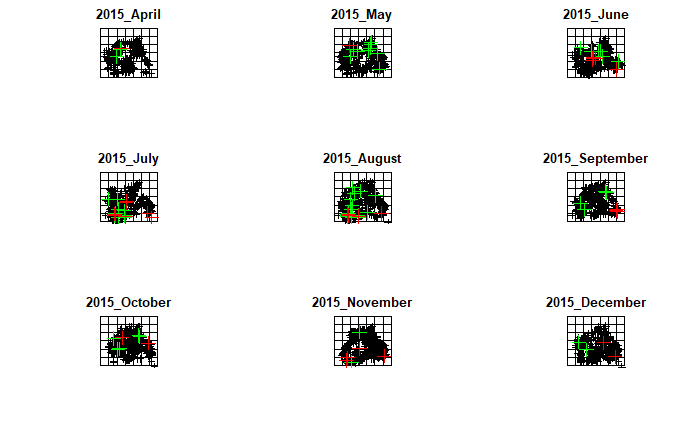

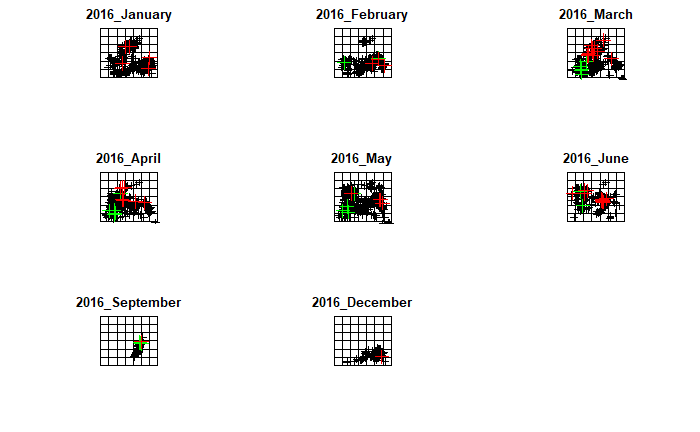

Here after we provide a monthly map with record of any GPS waypoints (in black), animal signs (in green) and poaching-related signs (in red) per sampling occasion in the SWS study area

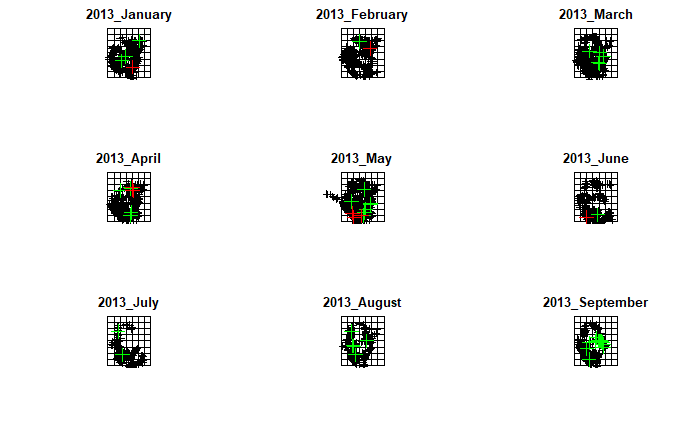

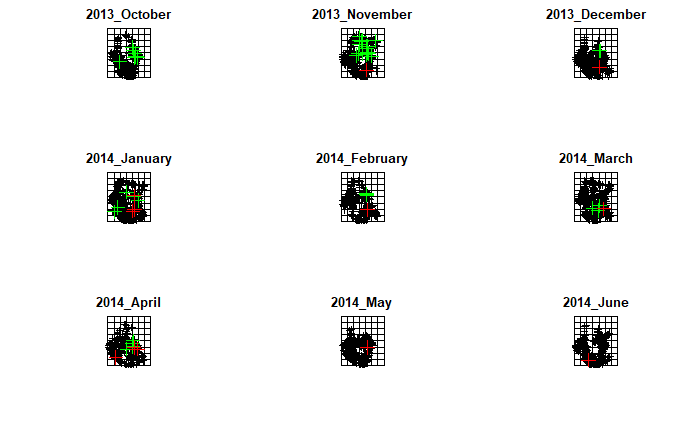

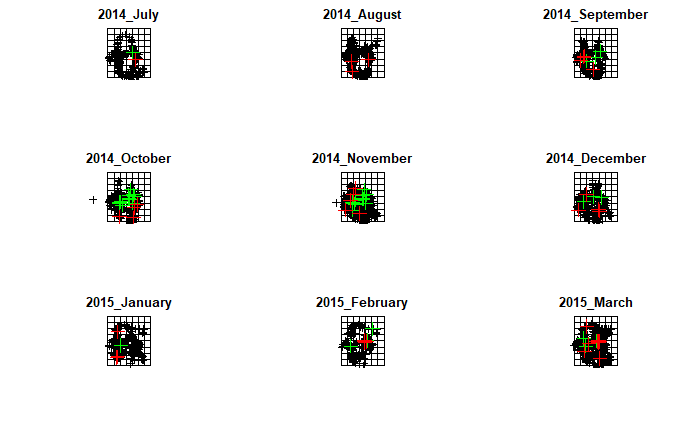

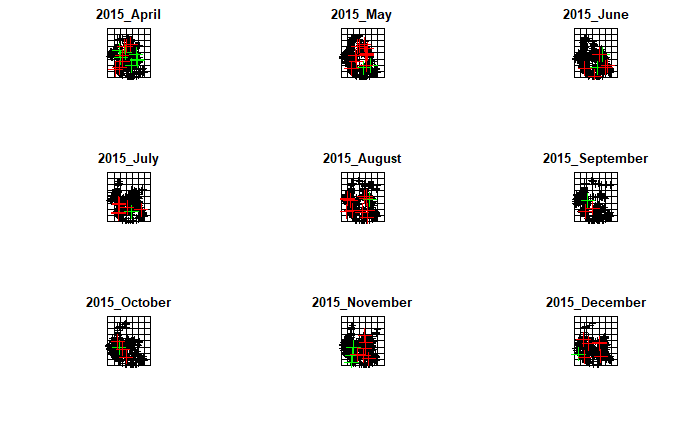

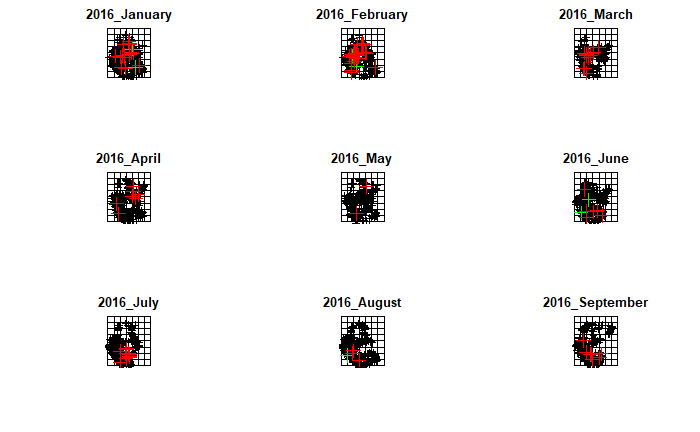

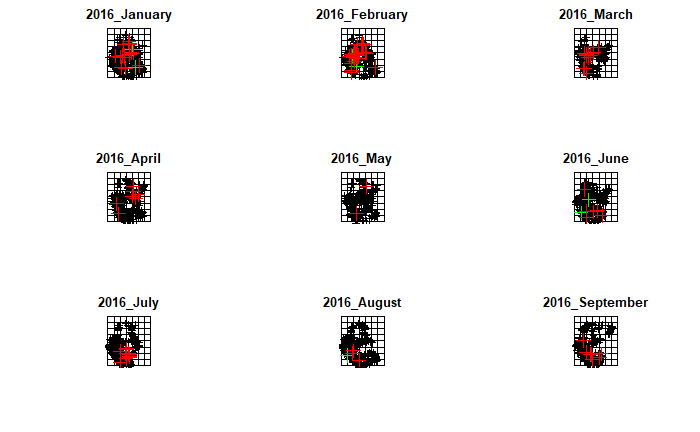

**Appendix C**

***SIMULATION STUDY***

We considered different scenarios first varying occupancy designs by the number of sites, number of repeated surveys (secondary occasions), and number of seasons (primary occasions), second by varying species detectability and spatial coverage then leading to more or less heterogeneous and sparse data sets. The more sites, the more heterogeneous the data can be. The higher the detection and occupancy probabilities the less sparse data is. We could then assess the bias that could emerge under all these different scenarios

**1 Methods**

*Occupancy design*

A key aspect of designing occupancy studies relies on the choice of the number and the size of sites as well as on the number of repeated surveys (i.e., the number of secondary occasions) that should be conducted and the length of the study period (i.e., the number of primary occasions) (MacKenzie & Royle 2005, Bailey et al. 2007). Here, we looked at fictive occupancy designs for analyzing co-occurrence of species, in our case study interactions between poachers and wildlife, in order to determine the most cost-effective allocation of ranger patrols. We did so in a similar way than MacKenzie and Royle (2005) simulation study aiming to optimize the sampling design for species distribution modelling. The authors made a general recommendation for managers to sample more sites with fewer surveys for the monitoring of rare species, and to sample few sites with more surveys for the survey of common species with a similar detection probability. Another study showed that both strategies are optimal for detecting a decline in species occupancy in the multi-season version of the model (Fields et al. 2005). To our knowledge, no study has yet investigated such questions in the framework of a two-species dynamic occupancy model.

In this first simulation study, we asked the question whether this general recommendation still holds when two species have different ecology, one with restricted range (wildlife) and the other with a more widespread distribution (poachers), assuming both were quite difficult to detect. The major goal of these simulations was to investigate, whether the occupancy design chosen was appropriate to our case study. We started by simulating similar study design than in our SMART data analysis, assuming 12 repeated surveys over 4 years across 81 sites. Across those fictive sites, we simulated a population of poachers with increasing occupancy range and a population of wildlife with decreasing occupancy range.

We simulated 1000 data sets akin to the SMART data describing the co-occurrence of wildlife and poachers based on the estimated parameters obtained from the best model of the joint analysis of the two study areas in Cambodia, the Serepok and Phnom Prich Wildlife sanctuaries (Table C.1). We then fitted the dynamic two-species occupancy model as presented in the methods and calculated the bias in parameter estimates. We used this simulation as a way to validate our empirical results obtained from the analysis of the SMART data on the impact of poaching in Cambodia.

We then calculated the bias and minimum squared error (MSE) for different occupancy design varying from few sites with many surveys to many surveys with few sampling sites and all other alternative combinations (many sites many surveys, few sites few surveys). We considered fictive study areas decomposed into 35, 100, 250 or 500 sites monitored during 3, 5 or 10 years (number of primary occasions) every month, trimester or semester (so with 3, 6 or 12 repeated surveys). For each combination of occupancy-design, we again simulated 1000 co-occurrence data sets based on the estimated parameters obtained from the best model of the joint analysis of the two study areas (Table C.1).

*Data sparseness*

In the second simulation study, we asked whether different level of sparseness in the data can generate some errors in parameter estimation which would need to be corrected when studying species, more or less difficult to detected and with more or less extended distribution range. We investigated the bias and MSE in all parameter estimates obtained after fitting the model to data sets simulated along a range of values for the occupancy and detection probabilities. We considered fictive populations for the two species from rare to abundant ($\psi_{P/W}, \psi_{W}, \psi_{P/\bar{W}}$ = {0.1, 0.25, 0.5}) from easily detectable to very discrete ($p_{P}=p_{W} =p_{WP}$ = {0.1, 0.25, 0.5}). For the sake of simplicity and computation time, we assumed detection probability was the same between poachers and wildlife. We also set the transition parameters constant and at equivalent values to the estimates obtained from the best model of the joint analysis (Table C.1). For each combination of species occupancy and detectability, we simulated 1000 occupancy datasets, designed with only 35 sites, with 12-repeated surveys, over 5 primary occasions. We chose a small number of sites to investigate what could be the bias in estimates if we analyzed the PPWS study area only.

**2. Results**

*Overview*

Results from the two simulation analyses revealed that bias tended to be more sensitive to occupancy design than to the parameters values used to simulate the data. Minimum squared error was consistently low across both simulation analyses (MSE < 10^-2^). Overall, our results showed that it is better to increase the number of repeated survey (secondary occasions) rather than increasing the number of sampled sites. When this condition was fulfilled, all parameter estimates were unbiased, except the occupancy probability of the poachers in absence of the wildlife with which they interact.

*Designing a two-species occupancy study*

In the first simulation analysis, aiming to explore the optimal occupancy design for two-species, we found that increasing survey effort by 4 (conducting 12 repeated survey instead of 3) decreased the mean bias by 20% whereas increasing the spatial coverage by 7 (sampling 250 sites rather than 35) decreased the mean bias by only 8%. Our results showed that the parameters most likely to be biased were the occupancy probabilities (more than 10% on average), and an underestimation in the probability that a site occupied by wildlife become extinct, but the biases diminished with increasing number of repeated surveys (Table C.2). The mean bias on occupancy estimates were 19% when only three secondary occasions were considered, 13% when six secondary occasions were recorded, down to only 3% in studies with 12 secondary occasions. The mean bias for any other estimate remains invariant to the number of sampled sites and was around 9%. Results from these simulations showed that the design we used for our analysis of poacher and wildlife interactions should not generate any bias greater than 15% (Table C.2).

*Sensitivity to occupancy and detection probabilities*

Results from the second simulation analysis with varying occupancy probabilities revealed that the biases in the estimates of all parameters were consistently lower than 5% and the associated MSE lower than 0.01 except for the occupancy probability of poachers in absence of wildlife (Figure C.1). This latter parameter was strongly underestimated (min(Bias$\psi_{P/\overline{W}}$) = - 0.3 MSE=0.01) when the probability of poachers occupancy in absence of wildlife was higher than in presence of them and overestimated (max(Bias$\psi_{P/\overline{W}}$) =0.3 MSE=0.002) when the occupancy probability of poachers was higher in presence of wildlife than in their absence (Figure C.1). However, the bias disappeared when occupancy probability of both poachers and wildlife were independent from each other, which was the case in our empirical analysis. This symmetrical trend observed in the bias of occupancy of poachers in absence of wildlife was consistent across the gradient of detection probabilities and probability of occupancy of wildlife (Figure C2.1).

*Discussion about potential effect of heterogeneity among sites*

In single species occupancy study under no financial constraint, the ultimate monitoring design is to sample many sites to reduce the sampling error with many survey and many sampling occasions to increase detection events. Our results from the first simulation show that the optimal design was not necessarily to maximize the number of sampling units even when there is a great number of sampling occasions (Table C2). We found that on average bias in parameter estimate increased by 1% when the number of sites increased from 100 sites to 500

Such result suggest that in the case of two-species occupancy designs where species have opposite range dynamics (one with a colonizing trend a extinction trend), increase the number of sampling units may generate more heterogeneity in the data and increase the bias. Increasing the sample size reduces sampling variability, which is, of course useful, but it does little to reduce concerns about unobserved bias (Rosenbaum 2005).

Our results suggest instead that the best design might be a trade-off between having sufficient number of repeated surveys to get robust estimates of detection and sufficient number of sites to minimize sampling variance for a rare species but not too many to avoid heterogeneous data. We acknowledged that this is only a preliminary simulation study which first aim was to validate our empirical analysis. We recommend further studies simulating other type of range dynamics and species interactions, and explicitly model the bias owed to detection heterogeneity by simulating data sets considering two classes of sites with different detection probabilities (Louvrier et al. 2019). Results from the second simulation study showed insignificant bias in the most homogeneous simulated data sets, those with the occupancy of a species independent of the occupancy of the other (Figure C.1). Here, we conclude that heterogeneity in detection and occupancy owed to the dependency of one species on the other may be an important source of bias in dynamic two-species occupancy model and we encourage further investigation on this topic.

Table C.1: Values used to simulate the data in the first part of the simulation study wherein we investigated the bias in parameter estimates emerging from different occupancy. The input parameter used to simulated the data were fixed to the values obtained from the empirical analysis.

| Symbol | Probability | Estimates from the best occupancy model |
| --- | --- | --- |
| $\Psi_{P/W}$ | Occupancy probability of poachers in presence of animals | 0.06 |
| $\Psi_{P/\bar{W}}$ | Occupancy probability of poachers in absence of animals | 0.06 |
| $\psi_{W}$ | Occupancy probability of animals | 0.46 |
| $\gamma_{W/\bar{P}}$ | probability that an unoccupied site is colonized by animals only | 0.28 |
| $\gamma_{P/\bar{W}}$ | probability that an unoccupied site is colonized by poachers only | 0.001 |
| $\gamma_{WP}$ | probability that an unoccupied site is colonized by both | 0.001 |
| $\gamma_{W/P}$ | probability that a site is colonized by animals given the presence of poachers | 0.05 |
| $\gamma_{P/W}$ | probability that a site is colonized by poachers given the presence of animals | 0.001 |
| $\epsilon_{W/P}$ | probability that animals are going extinct given the presence of poachers in a site | 0.80 |
| $\epsilon_{P/W}$ | probability that poachers are going extinct given the presence of animals in a site | 0.20 |
| $\epsilon_{WP}$ | the probability that poachers and animals are going extinct in a site | 0.001 |
| $\epsilon_{P/\bar{W}}$ | the extinction probability of poachers conditional on the absence of animals | 0.20 |
| $\epsilon_{W/\bar{P}}$ | the extinction probability of animals conditional on the absence of poachers | 0.41 |
| $\omega_{WP}$ | probability that a site occupied by animals is replaced by poachers | 0.44 |
| $\omega_{PW}$ | probability a site occupied by poachers only is replaced by animals | 0.12 |

Table C.2: Bias in the 18 parameter estimates of the two-species dynamic occupancy model calculated from 1000 simulations run for 36 different occupancy design : with number of sites ranging from 35, 100, 250, 500 monitored during 3, 5, 10 primary occasions with 3, 6, or 12 secondary occasions. We calculated the bias as the mean difference between the parameter values estimated and the values used to simulate the data across 1000 simulations. Yellow colored cells represent a positive bias greater than 0.1 and blue colored cell a negative bias lower than -0.1. The framed row corresponds to the bias in estimates for the occupancy design the closest to ours. The value used to simulate the data are presented in table 1 of the appendix (See table 1 for the signification of each parameter symbol).

| Site | Occasions | | Mean Bias in parameter estimates | | | | | | | | | | | | | | | | | |
| --- | --- | --- | --- | --- | --- | --- | --- | --- | --- | --- | --- | --- | --- | --- | --- | --- | --- | --- | --- | --- |
| # | 1ary | 2ary | $\psi_{w}$ | $\psi_{P/\bar{W}}$ | $\psi_{P/W}$ | $\gamma_{wP}$ | $\gamma_{P}$ | $\gamma_{W}$ | $\varepsilon_{P/\bar{W}}$ | ω_WP_ | $\gamma_{P/W}$ | $\varepsilon_{W/\bar{P}}$ | ω_PW_ | $\gamma_{W/P}$ | ε_WP_ | ε_W_ | ε_P_ | $p_{W}$ | $p_{P}$ | $p_{WP}$ |
| 35 | 3 | 3 | 0.18 | 0.12 | 0.12 | 0.21 | 0.20 | 0.03 | 0.09 | -0.04 | 0.07 | 0.08 | -0.07 | -0.01 | 0.53 | -0.59 | 0.01 | 0.03 | 0.03 | -0.02 |
| 100 | 3 | 3 | 0.27 | 0.31 | 0.18 | 0.25 | 0.08 | 0.06 | 0.02 | -0.03 | 0.06 | -0.09 | -0.02 | 0.10 | 0.46 | -0.49 | -0.04 | 0.01 | 0.01 | -0.01 |
| 250 | 3 | 3 | 0.12 | 0.08 | 0.10 | 0.05 | 0.01 | -0.01 | 0.00 | 0.01 | 0.01 | -0.10 | -0.01 | 0.09 | 0.29 | -0.34 | -0.01 | 0.00 | 0.01 | 0.00 |
| 500 | 3 | 3 | 0.21 | 0.17 | 0.20 | 0.29 | -0.05 | 0.02 | -0.01 | 0.01 | 0.01 | -0.27 | -0.01 | 0.27 | 0.38 | -0.37 | -0.05 | 0.00 | 0.00 | -0.03 |
| 35 | 5 | 3 | 0.27 | 0.19 | 0.22 | 0.10 | 0.22 | 0.03 | 0.02 | -0.02 | 0.09 | 0.00 | -0.05 | 0.03 | 0.42 | -0.59 | 0.03 | 0.02 | 0.01 | -0.02 |
| 100 | 5 | 3 | 0.17 | 0.17 | 0.11 | 0.11 | 0.05 | 0.01 | -0.02 | 0.00 | 0.05 | -0.09 | 0.00 | 0.08 | 0.33 | -0.45 | 0.01 | 0.00 | 0.01 | 0.00 |
| 250 | 5 | 3 | 0.25 | 0.25 | 0.21 | 0.30 | 0.03 | 0.04 | -0.04 | 0.02 | 0.03 | -0.21 | -0.02 | 0.22 | 0.37 | -0.46 | -0.05 | 0.00 | 0.00 | -0.03 |
| 500 | 5 | 3 | 0.06 | 0.04 | 0.07 | 0.03 | 0.00 | 0.00 | -0.02 | 0.00 | 0.01 | -0.10 | -0.01 | 0.10 | 0.13 | -0.17 | -0.01 | 0.00 | 0.00 | 0.00 |
| 35 | 10 | 3 | 0.32 | 0.18 | 0.25 | 0.21 | 0.05 | 0.06 | -0.04 | 0.02 | 0.06 | -0.03 | -0.03 | 0.06 | 0.45 | -0.54 | -0.01 | 0.01 | 0.01 | -0.02 |
| 100 | 10 | 3 | 0.29 | 0.26 | 0.22 | 0.18 | 0.19 | 0.02 | -0.06 | 0.03 | 0.05 | -0.19 | -0.02 | 0.20 | 0.25 | -0.48 | -0.04 | 0.00 | 0.00 | -0.02 |
| 250 | 10 | 3 | 0.11 | 0.10 | 0.10 | 0.03 | 0.05 | -0.01 | -0.03 | 0.01 | 0.02 | -0.12 | -0.01 | 0.12 | 0.14 | -0.23 | -0.02 | 0.00 | 0.00 | 0.00 |
| 500 | 10 | 3 | 0.27 | 0.32 | 0.24 | 0.10 | 0.16 | 0.04 | -0.06 | 0.02 | 0.04 | -0.25 | -0.02 | 0.26 | 0.19 | -0.37 | -0.05 | -0.01 | 0.00 | -0.03 |
| 35 | 3 | 6 | 0.11 | 0.12 | 0.03 | 0.06 | 0.06 | 0.02 | 0.04 | -0.02 | 0.01 | -0.04 | 0.00 | 0.03 | 0.29 | -0.38 | 0.03 | 0.00 | 0.00 | 0.05 |
| 100 | 3 | 6 | 0.08 | 0.05 | 0.06 | 0.05 | -0.01 | 0.00 | -0.01 | -0.01 | 0.02 | -0.07 | 0.00 | 0.07 | 0.22 | -0.25 | -0.01 | 0.00 | 0.00 | 0.00 |
| 250 | 3 | 6 | 0.11 | 0.12 | 0.09 | 0.05 | -0.01 | 0.00 | -0.03 | 0.02 | 0.00 | -0.12 | 0.00 | 0.12 | 0.19 | -0.18 | -0.04 | 0.00 | 0.00 | -0.01 |
| 500 | 3 | 6 | 0.06 | 0.05 | 0.06 | 0.01 | 0.01 | -0.01 | -0.04 | 0.02 | 0.00 | -0.06 | 0.00 | 0.06 | 0.07 | -0.10 | -0.01 | 0.00 | 0.00 | 0.00 |
| 35 | 5 | 6 | 0.16 | 0.14 | 0.04 | 0.05 | 0.08 | 0.02 | -0.02 | 0.02 | 0.02 | -0.03 | -0.02 | 0.04 | 0.23 | -0.40 | 0.04 | 0.00 | 0.00 | 0.09 |
| 100 | 5 | 6 | 0.21 | 0.24 | 0.12 | 0.02 | 0.13 | 0.02 | -0.04 | 0.03 | 0.01 | -0.12 | -0.02 | 0.13 | 0.14 | -0.32 | -0.03 | -0.01 | 0.00 | 0.00 |
| 250 | 5 | 6 | 0.15 | 0.17 | 0.12 | 0.00 | 0.11 | -0.02 | -0.05 | 0.04 | 0.01 | -0.09 | -0.02 | 0.10 | 0.06 | -0.22 | 0.00 | -0.01 | 0.00 | -0.01 |
| 500 | 5 | 6 | 0.24 | 0.33 | 0.17 | 0.02 | 0.12 | 0.04 | -0.04 | 0.04 | 0.01 | -0.16 | -0.02 | 0.16 | 0.09 | -0.26 | -0.03 | -0.01 | 0.00 | -0.02 |
| 35 | 10 | 6 | 0.17 | 0.11 | 0.12 | 0.02 | 0.07 | -0.01 | 0.04 | 0.00 | 0.03 | -0.03 | -0.05 | 0.07 | 0.18 | -0.41 | 0.03 | -0.01 | -0.02 | 0.01 |
| 100 | 10 | 6 | 0.13 | 0.12 | 0.12 | 0.01 | 0.10 | -0.01 | -0.01 | 0.02 | 0.02 | -0.08 | -0.03 | 0.10 | 0.11 | -0.28 | -0.02 | 0.00 | -0.01 | -0.01 |
| 250 | 10 | 6 | 0.16 | 0.17 | 0.12 | 0.01 | 0.13 | 0.00 | -0.04 | 0.03 | 0.02 | -0.10 | -0.02 | 0.11 | 0.07 | -0.23 | -0.01 | -0.01 | 0.00 | -0.01 |
| 500 | 10 | 6 | 0.14 | 0.19 | 0.10 | 0.01 | 0.13 | 0.00 | -0.03 | 0.02 | 0.02 | -0.08 | -0.02 | 0.08 | 0.07 | -0.19 | -0.02 | -0.01 | -0.01 | -0.02 |
| 35 | 3 | 12 | 0.04 | 0.02 | 0.01 | 0.02 | -0.02 | 0.01 | 0.00 | 0.00 | 0.01 | -0.04 | 0.03 | 0.02 | 0.13 | -0.20 | 0.02 | 0.00 | 0.00 | 0.10 |
| 100 | 3 | 12 | 0.09 | 0.07 | 0.05 | 0.01 | 0.04 | -0.01 | -0.02 | 0.02 | 0.00 | -0.07 | 0.00 | 0.06 | 0.11 | -0.15 | -0.01 | -0.01 | 0.00 | 0.00 |
| 250 | 3 | 12 | 0.08 | 0.02 | 0.08 | 0.00 | -0.04 | -0.03 | -0.04 | 0.02 | 0.01 | -0.07 | -0.01 | 0.07 | 0.03 | -0.12 | 0.05 | -0.01 | 0.00 | -0.02 |
| 500 | 3 | 12 | 0.10 | 0.04 | 0.07 | 0.00 | 0.01 | -0.02 | -0.02 | 0.01 | 0.01 | -0.06 | -0.01 | 0.06 | 0.05 | -0.12 | 0.02 | -0.01 | 0.00 | -0.01 |
| 35 | 5 | 12 | 0.02 | 0.01 | 0.01 | 0.01 | 0.01 | 0.00 | 0.03 | 0.01 | 0.02 | 0.03 | -0.02 | -0.02 | 0.20 | -0.36 | 0.06 | -0.01 | -0.02 | 0.22 |
| 100 | 5 | 12 | 0.07 | 0.04 | 0.05 | 0.00 | 0.01 | -0.01 | 0.00 | 0.02 | 0.01 | -0.01 | -0.02 | 0.02 | 0.09 | -0.22 | 0.02 | -0.01 | -0.01 | 0.05 |
| 250 | 5 | 12 | 0.08 | 0.06 | 0.08 | 0.01 | 0.05 | -0.01 | -0.03 | 0.02 | 0.01 | -0.06 | -0.02 | 0.07 | 0.04 | -0.15 | 0.01 | 0.00 | -0.01 | -0.01 |
| 500 | 5 | 12 | 0.11 | 0.11 | 0.06 | 0.00 | 0.05 | -0.02 | -0.03 | 0.03 | 0.01 | -0.04 | -0.01 | 0.04 | 0.05 | -0.23 | 0.06 | -0.01 | -0.01 | -0.02 |
| 35 | 10 | 12 | 0.10 | 0.06 | 0.04 | 0.01 | -0.07 | 0.01 | 0.14 | 0.07 | 0.02 | 0.04 | -0.08 | -0.08 | 0.24 | -0.53 | 0.10 | -0.01 | -0.05 | 0.41 |
| 100 | 10 | 12 | 0.01 | 0.06 | 0.16 | 0.00 | -0.14 | -0.04 | 0.38 | -0.03 | 0.04 | 0.12 | -0.18 | -0.05 | 0.22 | -0.57 | 0.10 | -0.06 | -0.11 | 0.35 |
| 250 | 10 | 12 | -0.32 | -0.03 | 0.07 | 0.00 | -0.17 | -0.04 | 0.76 | -0.15 | 0.01 | 0.52 | -0.37 | -0.15 | 0.28 | -0.58 | 0.12 | -0.10 | -0.16 | 0.40 |
| 500 | 10 | 12 | -0.36 | -0.05 | -0.03 | 0.00 | -0.17 | -0.04 | 0.80 | -0.17 | 0.00 | 0.55 | -0.38 | -0.16 | 0.31 | -0.58 | 0.10 | -0.10 | -0.16 | 0.31 |

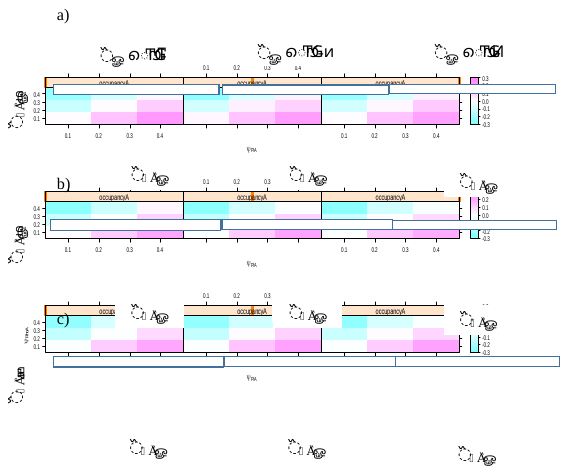

Figure C.1: Bias in the occupancy probability of poachers in absence of wildlife in relation to varying input values used to simulate data. We varied the occupancy probability of poachers in absence of wildlife (y-axis), the occupancy probability of poachers in presence of wildlife (x-axis), the occupancy probability of wildlife regardless of the occurrence of poachers (from the left to the right). The detection probability varied from a) 0.1, b) 0.25 to c) 0.4 and was assumed the same between poachers and wildlife.
